# The SoxC transcription factors regulate multiple early retinal lineages and function in parallel with Atoh7 to promote retinal ganglion cell genesis

**DOI:** 10.64898/2026.07.25.740745

**Authors:** Sandy Enriquez, Yichen Ge, Nan Nan, Véronique Lefebvre, Tao Liu, Xiuqian Mu

**Affiliations:** Department of Ophthalmology/Ross Eye Institute, Jacobs School of Medicine and Biomedical Sciences, University at Buffalo, Buffalo, New York; Department of Biochemistry, Jacobs School of Medicine and Biomedical Sciences, University at Buffalo, Buffalo, New York; Department of Biostatistics, School of Public Health and Health Professions, University at Buffalo, Buffalo, New York; Department of Surgery/Division of Orthopaedic Surgery, Children’s Hospital of Philadelphia, Philadelphia, Pennsylvania; Department of Biostatistics & Bioinformatics, Roswell Park Comprehensive Cancer Center, Buffalo, New York

**Author notes:** Corresponding author: Xiuqian Mu.

**Keywords:** Retinal development, Transcription factors, Cell lineage, Cell fate, Retinal ganglion cells, Gene regulation, DNA-protein interaction, Enhancers

## Abstract

Retinal development is orchestrated by a network of transcription factors that guide multipotent retinal progenitor cells (RPCs) to fate-committed lineages, ultimately producing seven major retinal cell classes. Among these, retinal ganglion cells (RGCs) serve as the sole output neurons of the retina, relaying visual and non-visual information to the brain. RGC specification requires a cascade of transcriptional regulators, including the basic helix-loop-helix (bHLH) factor Atoh7, which confers competence to RPCs, and Pou4f2 and Isl1, which drive terminal differentiation and subtype diversification. Previous studies demonstrate that the SoxC group transcription factors (Sox4, Sox11, and Sox12) are also involved in RGC genesis, but their precise integration into the Atoh7-driven regulatory hierarchy remains undefined. To address this question, we used the retina-specific Vsx2-Cre line to generate *Sox4/Sox11* double conditional knockout (dcKO) and *Sox4/Sox11/Atoh7* triple knockout (tKO) mice. Immunohistochemistry revealed profound lineage disruption and reduced progenitor proliferation and survival in both dcKO and tKO retinas; not only RGC genesis but also that of horizontal and amacrine (H&As) cells were severely compromised, whereas photoreceptor cells (PHCs) production increased. These results indicate that Sox4 and Sox11 are involved in the coordinated generation of the different early retinal lineages. Like the *Atoh7*-null retina, RGC precursors still formed in the SoxC dcKO retina, but their genesis was almost completely abolished in the tKO retina. Our findings indicate that the SoxC factors act in parallel with Atoh7 as a major upstream regulatory input to initiate RGC fate. Bulk RNA-seq revealed the SoxC-dependent transcriptional programs and signaling pathways and confirmed the lineage changes demonstrated by marker analysis. CUT&Tag analysis identified the genome-wide binding sites and thereby the target genes of Sox11, further illuminating the mechanisms underlying functions of the SoxC factors in multiple retinal cell states/types during development.

## Introduction

The central nervous system possesses the most complex cellular diversity with different classes and subclasses of neurons and glial cells. How these diverse cell types are generated from initially undifferentiated progenitor cells is still poorly understood. As part of the central nervous system, the vertebrate retina serves as a powerful model for understanding how progenitor cells give rise to multiple neuronal and glial cell types because of its accessibility for experimental studies and the availability of many genetic tools. During retinal development, different cell types are generated in a conserved temporal order but can be grouped into two overlapping waves, with retinal ganglion cells (RGCs), cones, horizontal cells, and amacrine cells differentiating in the first wave, followed by rods, bipolar cells, and Müller glia in the second wave ^1^. In the mouse, the first wave occurs before birth from embryonic day (E) 11 to postnatal day (P) 0, whereas the second wave happens mostly after birth until around P10 ^2^. All seven retinal cell types arise from the multipotent retinal progenitor cells (RPCs), which transition through distinct cell states, including a transitional RPC (tRPC) state, to individual lineages ^3–7^. These fate decisions are shaped by extrinsic factors such as the Notch and Shh pathways, together with the intrinsic function of cell fate specifying transcription factors^8–11^. Our current report is concerned with the cell types being generated in the first wave. For the RGC lineage, Atoh7 (Math5) is a key transcription factor that confers competence to RPCs, while downstream factors such as Pou4f2 (Brn3b) and Isl1 drive RGC differentiation and axonogenesis^12–14^. Additional factors, including the SoxC factors, Dlx1/Dlx2, and Pou3f1, also participate in RGC genesis ^15–20^. Further differentiation and subtype specification of RGCs are carried out by a myriad of downstream transcription factors, but only a few of them have been characterized ^21–23^. For horizontal and amacrine cells (H&As), a different set of transcription factors are involved, such that Foxn4 activates Ptf1a to promote horizontal and amacrine fates, Tfap2a/b supports their differentiation, whereas the Onecut factors direct Ptf1a-expressing precursors toward the horizontal fate^24–28^. Further downstream, Lhx1 (Lim1) and Prox1 more specifically drive horizontal cell differentiation ^29,30^. For photoreceptors (PHCs), Otx2 and Neurod1 induce Crx and Neurod4 to specify the photoreceptor fate. Most photoreceptors generated during the early wave are cones defined by Thrb and Rxrg, whereas later on, Nrl and Nr2e3 direct photoreceptor precursors toward rods^31^. The temporal order of retinal cell type generation is due to changes in competence of the RPCs. Although the underlying mechanisms are not yet fully understood, a number of transcription factors regulating early versus late competence, including Lhx2, Ikaros, Casz1, the Nfi factors, and Foxp1, have been identified ^4,32–35^.

Despite past progress, the central question regarding how the retinal cell fate choices are made from multipotent RPCs remains poorly understood. Single cell transcriptomics and epigenomics studies have offered significant insights. It has now become clear that all the cell lineage trajectories available during any developmental time start from naive proliferating RPCs (nRPCs) and go through a shared transitional state named transitional or neurogenic RPCs (tRPCs) before final differentiation^3,36^. nRPCs are in a high proliferative state, have high levels of Notch pathway activity, and express a group of transcription factors, including Sox2, Pax6, Rax, Lhx2, Vsx2, Nr2e1, and Six3/6, which are required to maintain their proliferative properties and differential potential ^3,33,37–40^. On the other hand, tRPCs are characterized as being ready to exit, or having already exited, the cell cycle, having the Notch signaling pathway turned off, and co-expressing transcription factors essential for the different cell lineages ^3,41^. Further, global transcription factor occupancy studies show that transcription factors for different lineages (e.g. Atoh7 for RGCs and Otx2 for PHCs) are not only co-expressed in tRPCs, but also co-bind to enhancers of many lineage specific genes ^36^. These findings prompted us to postulate that these transcription factors drive tRPCs to different fates via competition and cross-repression^3,36^. Although there is strong evidence supporting this idea, the nature of the competition and the number of transcription factors involved remain unknown. Atoh7 is a key transcription factor expressed in all early tRPCs that give rise to all the cell types generated in the early wave^3,13,42^. As mentioned above, in the RGC gene regulatory network, Atoh7 is a crucial regulator of RGC specification and is required to activate RGC lineage factor genes such *Isl1* and *Pou4f2* ^12,14,43–45^ Deletion of *Atoh7* disrupts the RGC lineage and ultimately results in the loss of almost all mature RGCs^12,43^. However, the RGC precursors are still generated in the *Atoh7*-null retina, and *Pou4f2* and *Isl1* are still activated, although at reduced levels ^3^. Nevertheless, these mutant RGC precursor cells do not fully differentiate and eventually die by apoptosis ^3,13^. Consistently, activities of many enhancers associated with RGC-specific genes are reduced, but substantial levels remain ^36^. Together, these observations suggest that additional factors cooperate with and act in parallel to Atoh7 in tRPCs to initiate the full RGC differentiation program. Identification of the additional factor(s) is necessary to further understand how the RGC lineage emerges among the different possible fate choices.

Sox4, Sox11, and Sox12 are members of the SoxC group of HMG-box transcription factors ^46^. They regulate cell fate and differentiation across multiple organs, including the central and peripheral nervous systems, heart, skeleton, and inner ear ^47–58^. Importantly, the SoxC factors are also involved in RGC formation. These factors function redundantly in the process, as knockout of individual genes leads to only moderate defects, but double knockout of *Sox4* and *Sox11* or triple knockout of *Sox4, Sox11* and *Sox12* severely compromises RGC development, disrupts optic nerve formation, and causes retinal hypoplasia^18,19,59^. In addition, loss of SoxC factors disrupts the formation of contralateral RGCs, which are 95% of RGCs in the mouse retina, while ipsilateral RGC differentiation is less affected^19^. Beyond development, the SoxC factors have been implicated in regenerative responses, where Sox11 overexpression can promote axon regrowth in select but not all RGC types ^60^. Together, these findings suggest that SoxC transcription factors play key regulatory roles in RGC genesis and function. Nonetheless, how SoxC factors promote the RGC lineage and how they intersect with Atoh7 and other transcription factors remain to be studied. Our scRNA-seq analysis suggests that the SoxC factors are expressed in multiple cell states/types, but highly in tRPCs and RGCs, and their expression is not affected in the *Atoh7*-null retina ^3^. We thus hypothesize that the SoxC factors function in parallel to Atoh7 to promote RGC fate in tRPCs.

In this report, we investigate the roles of the SoxC factors in driving tRPCs toward the different early retinal lineages, particularly RGCs. Although all three SoxC factors are expressed in the developing retina and function redundantly, Sox4 and Sox11 appear to play the major roles, as Sox12 is expressed at relatively low levels^18,46^. Therefore, we focus on Sox4 and Sox11 in this study. Because of their broad roles, germline deletion in the mouse of Sox4 or Sox11 is embryonic lethal ^61,62^. To elucidate how Sox4 and Sox11 contribute to RGC genesis, we generated *Sox4/Sox11* double conditional knockout (dcKO) and *Sox4/Sox11/Atoh7* triple knockout (tKO) embryos/mice and examined the cellular phenotypes by immunohistochemistry (IHC) and transcriptomic changes by bulk RNA-seq. Our results indicate that, similar to the *Atoh7*-null retina, RGC precursor cells still form in the dcKO retina but fail to further differentiate and eventually die. In the tKO retina, however, RGC precursors fail to form, suggesting that the SoxC factors and Atoh7 indeed work together to promote the RGC lineage. Interestingly, horizontal and amacrine populations also decline, whereas photoreceptors expand, in the dcKO and tKO retinas. Reduced proliferation and increased apoptosis were also evident in both dcKO and tKO retinas. To further understand the mechanisms underlying the function of Sox4 and Sox11, we also performed CUT&Tag to identify genome-wide binding sites of Sox11 and thereby their direct target genes. Our results not only confirm that the SoxC factors function in parallel with Atoh7 to promote the RGC lineage but also suggest that they function in multiple cell states to regulate nRPC proliferation and coordinate production of the multiple early lineages from tRPCs.

## Materials and Methods

### Animals

All mice were maintained in the C57BL/6J genetic background. The floxed *Sox4* and *Sox11* alleles (*Sox4^F^*and *Sox11^F^* respectively) and the Vsx2-Cre (*Vsx2^Cre^*, also known as Chx10-Cre) allele were all described previously ^61,63,64^. Double conditional knockout (dcKO, *Sox4^F/F^;Sox11^F/F^;Vsx2^Cre^*) mice delete *Sox4* and *Sox11* in retinal progenitor cells from E10.5 ^63^, and littermate controls, *Sox4^F/F^;Sox11^F/F^*, are phenotypically wild type. The *Atoh7^lacZ^* allele, which has been described previously, was used as an *Atoh7* null allele ^12^. Triple knockout (tKO) mice (*Sox4^F/F^;Sox11^F/F^;Vsx2^Cre^;Atoh7^LacZ/LacZ^*) lack all three genes in the retina, and the corresponding controls, *Sox4^F/F^;Sox11^F/F^;Atoh7^LacZ/+^,* are phenotypically normal. Breeder mice were 8 weeks to 6 months of age. Efforts were made to use littermate control and knockout embryos and pups, but due to the need to cross multiple alleles, those from age-matched litters were also used. In either case, multiple litters for each genotype were used to ensure reproducibility. Wild type C57BL/6J mice of the same age range were also bred to obtain embryos and pups for RNAscope in situ hybridization, immunohistochemistry, and other histological assays. Mice of both sexes were used for all experiments, and no overt sex differences were observed. All procedures using mice conformed to the U.S. Public Health Service Policy on Humane Care and Use of Laboratory Animals and were approved by the Institutional Animal Care and Use Committees of Roswell Park Comprehensive Cancer Center and the University at Buffalo.

### Immunofluorescence

Immunofluorescence staining was performed as described previously ^3,65^. Timed matings were established, and embryos were collected at the desired stages for immunofluorescence on retinal cryosections. Embryos (E14.5 or earlier) or embryo heads (E17.5 or later) were fixed in 4% paraformaldehyde for 0.5 hrs and samples were rinsed in PBST (PBS, pH 7.4, with 0.2% Tween-20) and cryoprotected through a sucrose gradient (10% then 20% sucrose for 1–1.5 hrs each, followed by 30% sucrose overnight). Tissue was embedded in OCT (Sakura Finetek USA, Inc.) and sectioned at 16 µm. Sections were washed with PBST (3 x 10 mins) and blocked for 1 hr with AdvanBlock-Chemi blocking solution (Advansta, Inc.). Following blocking, primary antibodies were applied overnight at 4 °C, followed by secondary antibody incubation the next day after PBST washes. Nuclei were counterstained with Hoechst 33342, and coverslips were mounted using AquaMount (Lerner Laboratories). Four central sections from two different embryos were used for each marker, and positive cells were counted manually on a per section basis. Significance of differences were determined by two-tailed, two-sample equal variance student t test.

The following primary antibodies were used: guinea pig anti-Sox4 ^66^ (1:1500, RRID: AB_2722602, gift from Dr. Elisabeth Sock), guinea pig anti-Sox11 ^66^ (1:2000, RRID: AB_2722601, gift from Dr. Elisabeth Sock), rabbit anti-Sox11 (1:1000, HPA000536, Sigma), goat anti-Pou4f2/Brn3b (1:100, SC-6026, Santa Cruz Biotechnology), mouse anti-Pou4f1/Brn3a (1:200, MAB1585, Millipore), goat anti-Isl1 (1:100, AF1837, R&D Systems), rabbit anti-Ebf1 (1:500, AB10523, Millipore), mouse anti-Tfap2a (1:200, 3B5, DSHB), rabbit anti-Ptf1a/P48^25^ (1:800, gift from Dr. Helena Edlund), rabbit anti-Crx (1:200, PA5-111077, Invitrogen), rat anti-Prdm1 (Blimp1) (1:100, sc-47732, Santa Cruz Biotechnology), mouse anti-Ki67 (1:200, 550609, BD Biosciences), and mouse anti-phospho-histone H3 (pH3; 1:50, 9706, Cell Signaling Technology). Species-appropriate Alexa Fluor 488– and Alexa Fluor 546–conjugated secondary antibodies from Thermo Fisher Scientific and Abcam were used for the primary antibodies. Images were taken using the Leica SP8 STED Confocal Microscope with the same settings for the experimental and control samples. Occasionally, image contrast was adjusted with Photoshop, but control and experimental images were adjusted to the same degree.

### RNAscope

RNAscope in situ hybridization was performed using the RNAscope 2.5 HD Assay RED kit (Advanced Cell Diagnostics) on both paraffin-embedded and pre-fixed OCT-embedded retinal tissues. For paraffin samples, embryos or embryonic heads were fixed in 4% paraformaldehyde (PFA) in PBS overnight, dehydrated, embedded in paraffin, and sectioned at 7 µm. For OCT samples, embryos or embryonic heads were processed as described above and sectioned at 16 µm. Sections were processed following the manufacturer’s protocols for target retrieval, and then hybridized with RNAscope probes against mouse *Sox4*, *Sox11*, and *Atoh7* (Advanced Cell Diagnostics). Signal amplification and detection were carried out using the RED chromogenic substrate supplied in the kit, followed by hematoxylin counterstaining, air-drying, and cover slipping. Images were taken using a Nikon Eclipse 80i microscope.

### TUNEL and EdU labeling

Apoptotic cells were detected using the ApopTag Plus In-Situ Apoptosis Fluorescein Detection Kit (S7111, Sigma-Aldrich) on OCT-embedded retinal sections. Sections were processed according to the manufacturer’s instructions. Briefly, sections were equilibrated in ApopTag Equilibration Buffer, incubated with the terminal deoxynucleotidyl transferase (TdT) reaction mixture to label DNA strand breaks, followed by incubation with fluorescein-conjugated anti-digoxigenin antibody. After washing, nuclei were counterstained with Hoechst (1:2000), cover slipped with aqueous mounting medium and imaged by confocal microscopy (Leica SP8 STED).

S phase retinal cells were labeled in vivo using the Click-iT EdU Cell Proliferation Kit for Imaging, Alexa Fluor 555 dye (Thermo Fisher Scientific), following the manufacturer’s instructions. Pregnant female mice carrying embryos at the indicated developmental stages received an intraperitoneal injection (IP) of EdU (25ug/g). After a 90 min incorporation period, dams were sacrificed, embryos were collected, and eyes were dissected. Embryonic eyes were fixed in 4% paraformaldehyde (PFA) in PBS, cryoprotected in sucrose, embedded in OCT, and cryosectioned. EdU detection on retinal sections was performed using the Click-iT reaction cocktail according to the kit protocol, and nuclei were counterstained with Hoechst (1:2000) before mounting and fluorescence imaging.

### Bulk RNA sequencing and analysis

For bulk RNA-seq, timed matings were set up, and E14.5 and E17.5 embryos were collected. Retinas were dissected and immediately stored in RNAlater (Invitrogen) while genotyping was performed, and stored at −80 °C. Retinas were pooled by genotype and developmental stage to generate four independent biological replicates for each genotype. However, one E14.5 dcKO replicate and one E14.5 tKO replicate were eliminated in the analysis due to incomplete Cre-mediated gene deletion. Total RNA was isolated from pooled retinas using Qiagen’s miRNeasy Mini Kit according to the manufacturer’s instructions, and RNA integrity was verified by a Bioanalyzer before library preparation. RNA-seq libraries were generated using TruSeq Stranded RNA Library Preparation kit (Illumina) and sequenced on a NovaSeq 6000 sequencer with S2 flow cells (PE100) to obtain 100 bp paired-end reads.

Raw FASTQ files were processed on the University at Buffalo Center for Computational Research (CCR) high-performance computing cluster using a Snakemake-based pipeline (Snakefile, config.yaml, slurm.sh, and associated utility scripts), which is available at https://github.com/macs3-project/genomics-analysis-pipelines/. After initial quality control with FastQC and MultiQC, reads were aligned to the mouse reference genome (mm10) using STAR ^67^, and gene-level counts were obtained with featureCounts/htseq-count. The count data were then analyzed using DESeq2 and genes with very low counts were filtered automatically through the program’s independent filtering procedure. Normalization was performed using the median-of-ratios method to account for sequencing depth and RNA composition. Differential expression analysis was performed in R using DESeq2 for each contrast ^68,69^. Genes with a Benjamini–Hochberg adjusted p-value (adjp) < 0.05 and an absolute log2 fold change of ≥ 0.58 (≈1.5-fold change) were considered differentially expressed. Downstream visualization of DEGs was carried out in R, using pheatmap/ggplot2 for heatmaps and clustering, and EnhancedVolcano for volcano plots ^70–72^. Gene ontology and pathway enrichment analyses were performed with web-based tools including ShinyGO and String (GO and KEGG) ^73–75^. BioVenn was used for DEG overlap analysis ^76^.

### CUT&Tag experiment and data analysis

CUT&Tag was performed with E14.5 wild-type retinal tissues. After timed mating, E14.5 retinas were collected and dissociated into single cell suspension, from which nuclei were isolated. CUT&Tag experiments followed our previously published protocols using Epicipher’s CUT&tag kit with some modifications ^36,77^. The major modification was that after tagmentation by pAG-Tn5, the DNA was purified by phenol/chloroform extraction and alcohol precipitation, instead of using magnetic beads. We found that this modification resulted in more consistent DNA yields. We tried two anti-Sox11 antibodies and one anti-Sox4 antibody, but only the rabbit anti-Sox11 antibody (HPA000536, Sigma) yielded convincing results. The anti-H3K27ac antibody was from Active Motif (39133, RRID:AB_2561016). The analysis was performed as described previously using MACS2 Peakcall version MACS/2.2.7.1 ^78^. Sequence reads were aligned to the mm10 mouse reference genome using bwa-mem. Peaks were called under pair-end mode, and the minimum FDR cutoff threshold was set to 0.01. Two independent replicates for each factor were first analyzed by MACS2, and correlation of the two replicates was calculated using ucsc-bigwigCorrelate tool to ensure reproducibility, but the eventual peak calling was performed with pooled sequence reads from both replicates to increase signal-to-noise ratios. Visualization of peaks was performed using Integrative Genomics Viewer (IGV) or UCSC genome browser with bigwig files.

### Data deposition

The E14.5 and E17.5 bulk RNA-seq library sequence reads, and the Sox11 and H3k27ac CUT&Tag library sequence reads have all been deposited into the NCBI under the BioProject ID PRJNA1490285.

## Results

### Sox4/11 and Atoh7 are independent of each other in expression during retinal development

Previous reports have differed on which retinal populations express SoxC factors; some suggest they are expressed only in RGCs, whereas others indicate they have broader expression patterns^18,19,45,79^. These discrepancies may have resulted from the different sensitivities of the assays used. Based on our scRNA-seq analysis, at E13.5, *Sox4* and *Sox11* have similar expression profiles, which begin in proliferating naïve RPCs at low levels, reach high levels in tRPCs and RGCs during the peak window of RGC genesis ^3^. They are also expressed in the other cell lineages, high in H&As but low in PHCs. Further, although Sox4 and Sox11 are expressed in RGCs, our bulk RNA-seq and scRNA-seq studies show that the overall *Sox4* and *Sox11* levels are not affected in the *Atoh7*-null retina in which most RGCs are lost ^3^. To validate these findings, we performed RNAscope in situ hybridization for *Sox4* and *Sox11* in control and *Atoh7*-null retinas. At E14.5, both genes were robustly expressed in the neuroblast layer (NBL) and ganglion cell layer (GCL) in the wild-type retina, which was consistent with our scRNA-seq data (**Fig. 1A**). Importantly, and again in agreement with our transcriptomics studies, the expression of both genes and their proteins in the NBL did not change substantially in E14.5 *Atoh7*-null retina, although their expression in the GCL was missing due to the absence of RGCs (**Fig. 1A, Suppl. Fig. 1**). These findings support our hypothesis that the SoxC factors function independently of Atoh7 during RGC genesis. We further confirmed a previous report ^59^ that, at later stages, the expression patterns of the two genes diverged in the wild-type retina. At E17.5, whereas both were continuously expressed in progenitor cells and RGCs at low levels, Sox4 demonstrated higher expression in the developing amacrine cells than Sox11 **(Suppl Fig. 2**). At P0 and P5, although expression of both genes were substantially downregulated in RPCs and RGCs, *Sox4* continued to be expressed at a high level in the amacrine cells, whereas Sox11 only demonstrated very sparse signals (**Suppl. Fig. 2**).

**Figure 1.**
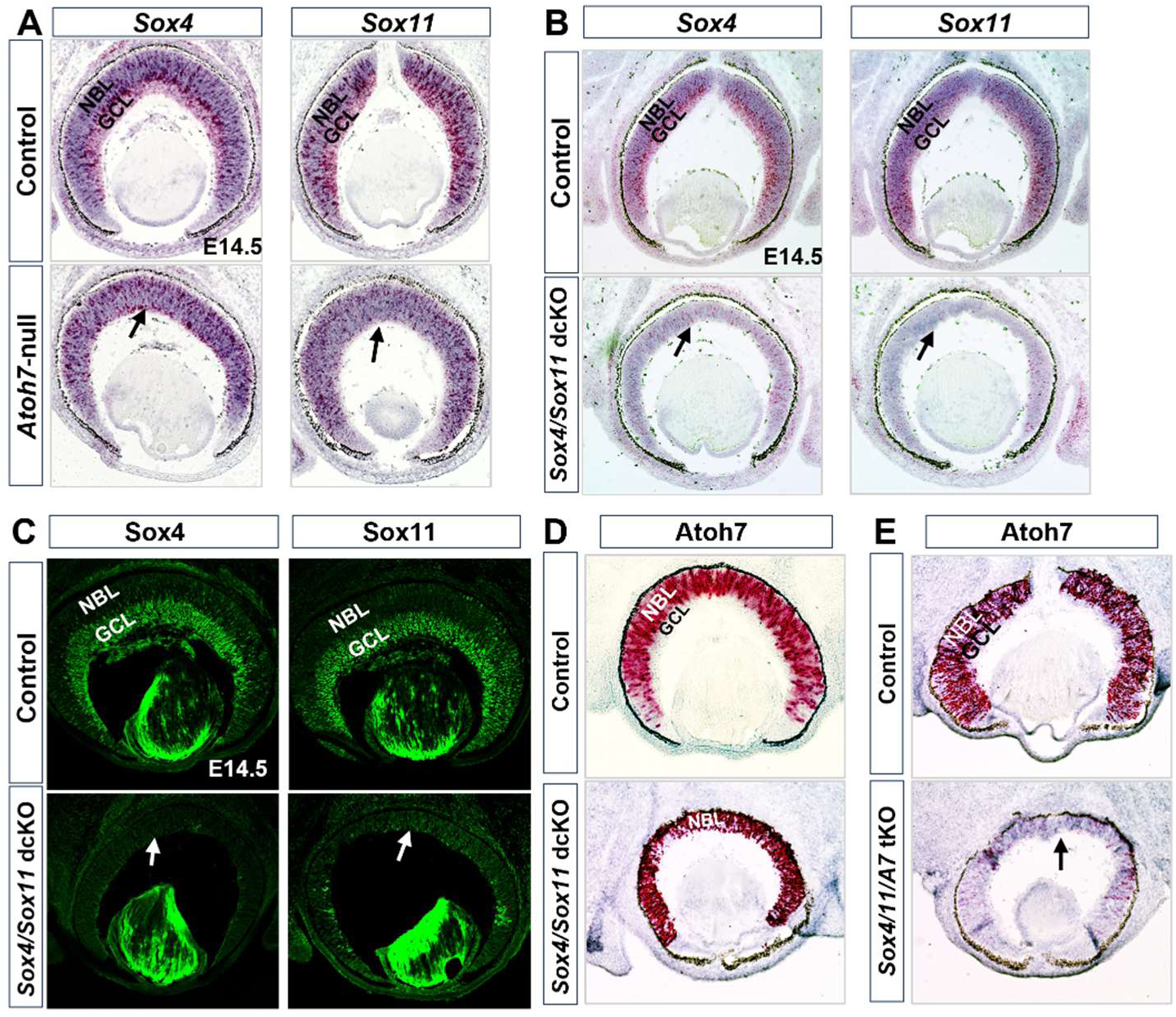
Independent expression of the SoxC factor genes and Atoh7 in the developing retina. **A.** RNAscope in situ hybridization of *Sox4* and *Sox11* in E14.5 wild type control (top) and *Atoh7*-null retinas (bottom) GCL: ganglion cell layer; NBL: neuroblast layer. Arrows point to the largely missing GCL in the mutant retina. **B.** RNAscope in situ hybridization indicates expression of *Sox4* and *Sox11* is abolished in the E14.5 *Sox4*/*Sox11* dcKO retina (bottom) in contrast to wild type control (top). **C.** Immunofluorescence demonstrates high Sox4 expression (top left) and Sox11 expression (top right) in GCL and low-level expression in the NBL of the E14.5 wild type control retina. (Bottom) Near complete loss of Sox4 and Sox11 expression in the dcKO retina. **D.** RNAscope in situ hybridization show Atoh7 is expressed in comparable levels in wild type control and dcKO retinas. E. Expression of Atoh7 is abolished in the *Sox4*/*Sox11*/Atoh7 tKO retina as compared to the wild type control. Arrows in the mutant panels point to the missing GCL.

We next bred the floxed *Sox4* and *Sox11* alleles (*Sox4^F/F^* and *Sox11^F/^*^F^) together and crossed with the *Vsx2^Cre^* line to delete these two genes (*Sox4^F/F^; Sox11^F/F^;Vsx2^Cre^*, dcKO) in the developing retina. To delete both SoxC genes and *Atoh7*, *Sox4^F/F^;Sox11^F/F^;Vsx2^Cre^* mice were crossed with *Atoh7^LacZ^* mice to generate *Sox4/Sox11/Atoh7* triple KO (*Sox4^F/F^;Sox11^F/F^;Vsx2^Cre^;Atoh7^LacZ/LacZ^*, tKO) retinas. The dcKO and tKO mice were viable but were smaller and less fertile than controls. RNAscope in situ hybridization demonstrated that both *Sox4* and *Sox11* mRNA were uniformly diminished in the E14.5 dcKO retina (**Fig. 1B**). Consistently, immunostaining with anti-Sox4 and anti-Sox11 antibodies showed near-complete loss of both proteins in the E14.5 dcKO retina (**Fig. 1C**). These results indicated that the two SoxC genes were both efficiently deleted in the retina. Importantly, *Atoh7* remained highly expressed in the dcKO retina as compared to the control (**Fig. 1D**) at E14.5, confirming that *Atoh7* was not dependent on the SoxC factors either ^18^. As expected, no *Atoh7* mRNA was detected in the tKO retina (**Fig. 1E**).

Both the dcKO (**Fig. 1B-D**) and tKO (**Fig. 1E**) retinas were thinner as compared to the wild-type control retinas and the GCL appeared to be largely diminished. The tKO retina, but not the dcKO retina, also manifested a coloboma phenotype, which persisted to at least postnatal day 0 (P0) (**Suppl. Fig. 3**). The postnatal dcKO and tKO eyes were smaller than the controls, and both lacked optic nerves, indicating defects in RGCs. Consistently, histological examination indicated that, at P5, both mutant retinas lacked the GCL, were much thinner, and the ONL did not demarcate from INL as compared to the controls (**Suppl. Fig. 4A, B**). At P22, the dcKO (**Suppl. Fig. 4A**) and tKO (**Suppl. Fig. 4B**) retinas showed severe hypoplasia and dysplasia, but the ONL and INL were discernible. The postnatal hypoplasia and dysplasia were at least partially due to secondary effects from the early developmental defects. Nevertheless, *Sox4* and *Sox11* may also play specific roles postnatally, since they are also expressed in the postnatal retina (**Suppl. Fig. 2**), but those roles could not be discerned in the current study using the *Vsx2^Cre^* line.

### Defective RGC genesis in *Sox4/Sox11* dcKO retina

Previous studies using different Cre lines suggested defective RGC genesis in the absence of *Sox4/Sox11* or *Sox4/Sox11/Sox12*, whereas single gene knockouts led to only modest defects ^18,20,59^. We first sought to validate those findings in the *Vsx2^Cre^* mediated *Sox4/Sox11* dcKO retina. Consistent with previous reports, immunostaining showed that the E14.5 dcKO retina exhibited a significant reduction of RGCs as marked by Pou4f2 and Isl1, two factors required for RGC differentiation and expressed in both nascent and differentiating RGCs, and Pou4f1 and Ebf1, two markers expressed only in differentiating RGCs already migrated to the forming GCL (**Fig. 2A**). Pou4f2+ and Isl1+ RGCs decreased by 66.2% and 69.0%, respectively, in the dcKO retina (**Fig. 2A and A’**). Notably, most remaining Pou4f2+ and Isl1+ RGCs in the dcKO retina were found in the NBL, and very few were observed in the GCL, indicating they were nascent RGC precursors (**Fig. 2A**). Consistently, Pou4f1+ and Ebf1+ RGCs, which normally have already migrated to the GCL, were reduced by 65.4% and 72.9%, respectively, in the dcKO retina (**Fig. 2A. and A’**). At E17.5, similar RGC loss persisted in the dcKO as revealed by the reduction of the four RGC markers. Overall, there was a 59.2% loss of Pou4f2+ and 64.3% percent reduction of Isl1+ cells (**Fig 2B and B’**). Most of the remaining Pou4f2+ and Isl1+ cells continued to be located largely in the NBL, indicating they were nascent RGC precursors. Very few Pou4f2+ or Isl+ RGCs were present in the GCL. Consistently, substantially fewer RGCs marked by Pou4f1 and Ebf1 were detectable in E17.5 dcKO retina, with 81.8% reduction of Pou4f1+ cells and 64.8% reduction of Ebf1+ cells (**Fig. 2B and B’**). Noticeably, there were more nascent RGC precursors in the dcKO retina as compared to the control at both E14.5 (**Fig. 2A**) and E17.5 (**Fig. 2B**), as indicated by Pou4f2 and Isl1. These increased cells were either due to compromised migration to the GCL, or more likely, to more RGC precursors being produced via the negative feedback mechanism during RGC production ^80–82^. A distinct row of Isl1+ cells was observed on the innermost side of the E17.5 dcKO retina (**Fig. 2B**). These were likely starburst amacrine cells, which normally reside right next to the GCL on the apical side ^83,84^.

**Figure 2.**
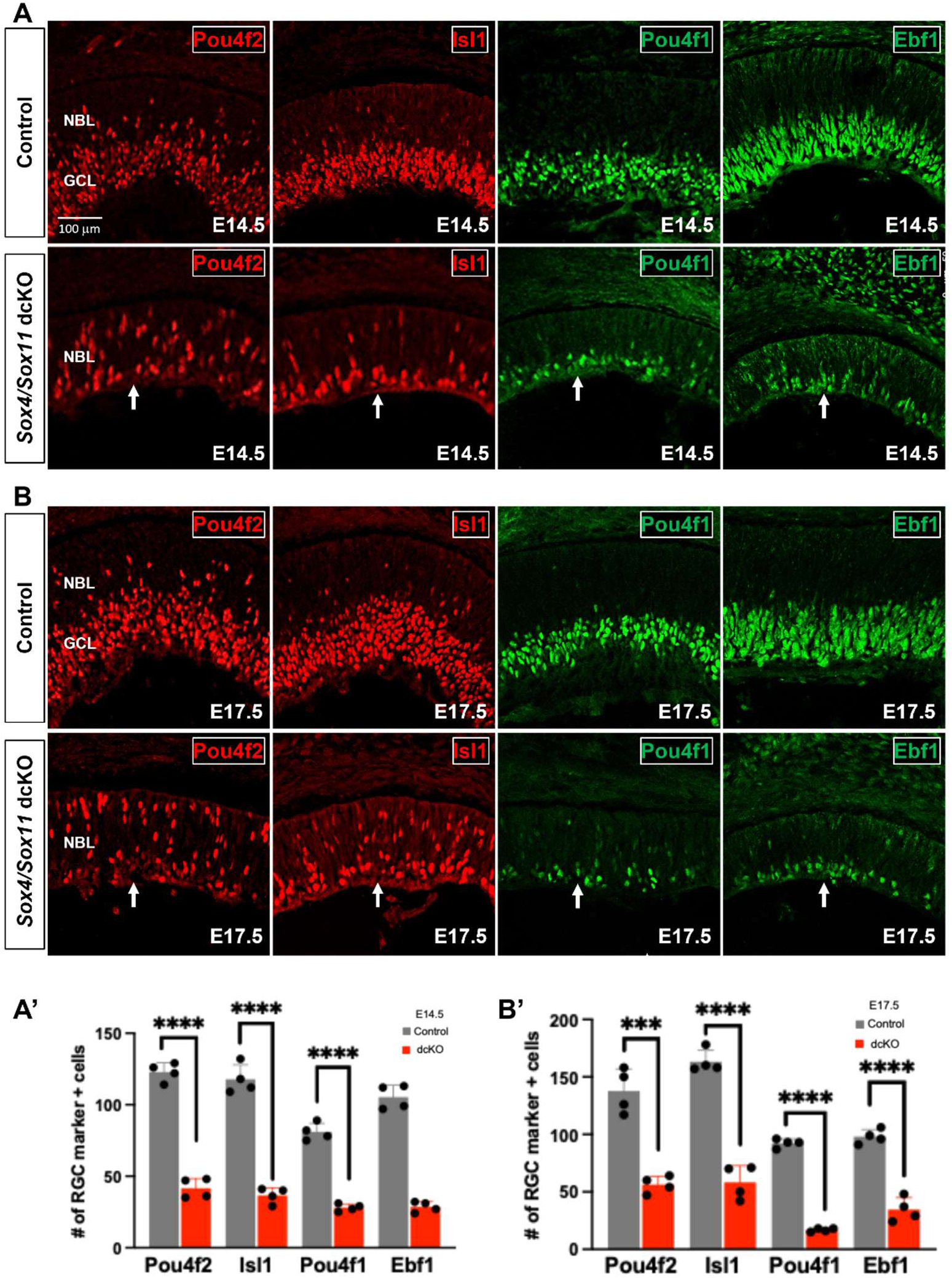
Deletion of Sox4 and Sox11 disrupts RGC development. **A.** Immunofluorescence staining of RGC markers in the E14.5 wild type (top) and dcKO (bottom) retinas. The GCL is much thinner in the dcKO as indicated by expression of all four markers, but nascent precursor RGCs marked by Pou4f2 and Isl1 are still present. **B.** Staining of the four RGC markers in the E17.5 retina further demonstrates the loss of the GCL but presence of RGC precursors in the NBL. **A’.** Quantification marker positive RGCs in **A**. **B’.** Quantification of marker positive RGCs in **B**. Error bars represent S.D. **** p<0.00001, *** p<0.0001. Arrows in **A and B** point to the thin GCL in the dcKOs. Scale bar applies to all panels in **A** and **B**.

These findings confirmed that the SoxC factors were indeed involved in RGC genesis. Importantly and not reported previously, RGC precursor cells still formed in the absence of Sox4 and Sox11, an observation similar to what occurs in the *Atoh7*-null retina as indicated by previous studies ^3^ and confirmed by immunostaining for Pou4f2 and Isl1 (**Suppl. Fig. 5**). These observations indicate that the SoxC factors and Atoh7 promote RGC lineage formation independently, but insufficiently. Like in the *Atoh7*-null retina, these precursor RGCs in the dcKO retina were lost eventually, as indicated by the reduced number of RGCs present in the GCL (**Fig. 2A, B**).

### SoxC transcription factors also regulate other early retinal lineages

The SoxC factors may also regulate the other early retinal lineages ^18^, but this has not been thoroughly investigated. In the time window of the first wave of retinal differentiation, cones, horizontal cells, and amacrine cells are generated concurrently with RGCs ^85^. Immunostaining for Ptf1a and Tfap2a, two transcription factors regulating horizontal and amacrine cell (H&As) differentiation ^25,27,86^, revealed severe defects in the formation of these cell types. At E14.5, substantially fewer Ptf1a+ cells and Tfap2a+ cells were present in the dcKO retina (**Fig 3A**). The Ptf1a+ cells were reduced by 67.8% and Tfap2a+ cells by 91.3% (**Fig. 3A’**). At E17.5, the reduction of H&As persisted, but to a lesser degree. Ptf1a+ precursor cells were reduced by 51.5% in the dcKO retina (**Fig. 3A and A’**). At this point, amacrine cells and horizontal cells have been demarcated to different positions, with amacrine cells occupying the inner side of the NBL next to the RGCs, and horizontal cells positioned in the middle region of the NBL, both positive for Tfap2a ^28^. This allowed us to determine that amacrine Tfap2a+ cells (**Fig. 3A, In NBL**) were reduced by 73.9% and horizontal Tfap2a+ cells (**Fig. 3A, Out NBL**) by 75.0% (**Fig. 3A’**).

**Figure 3.**
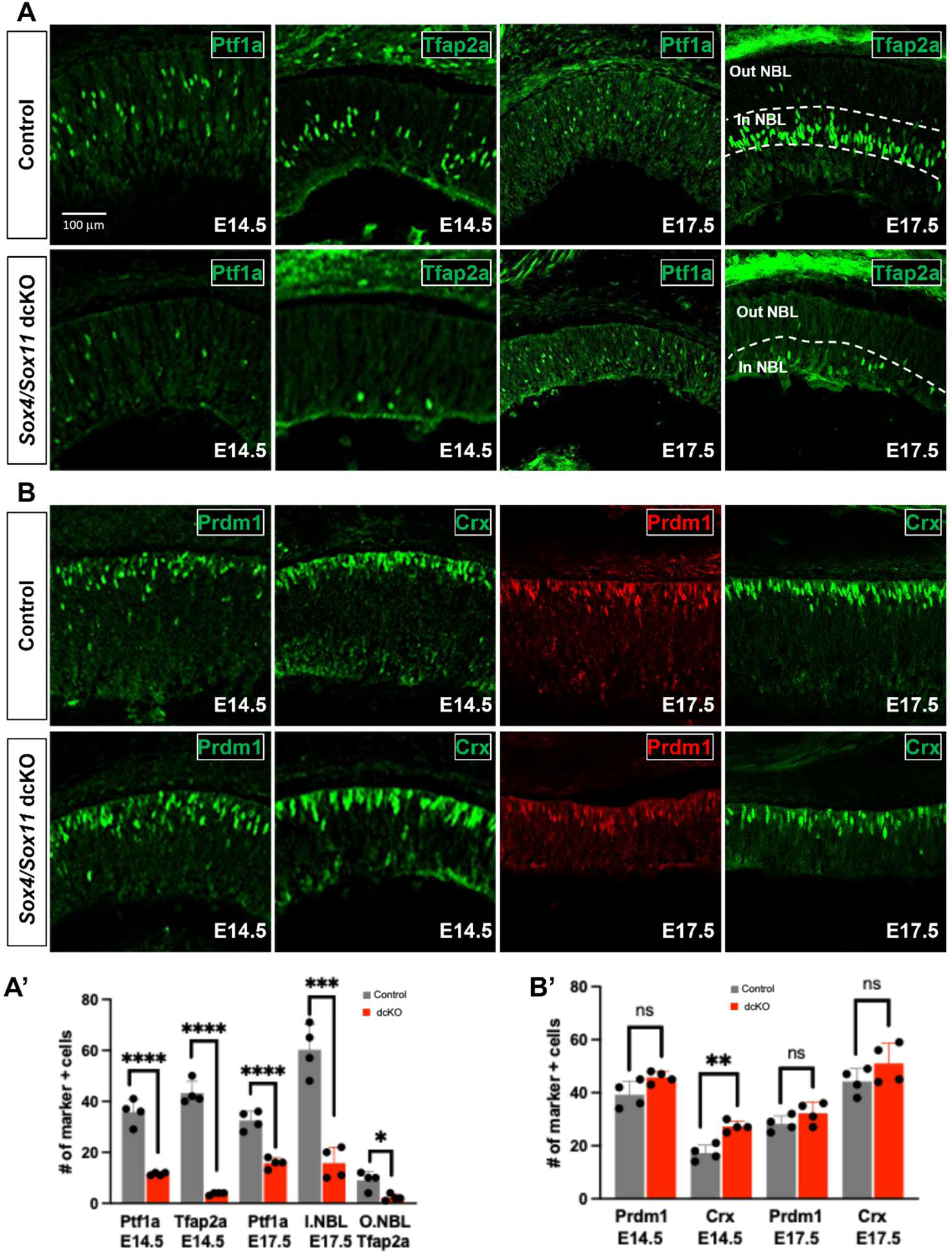
Decrease in H&As and increase in PHCs in the dcKO retina **A.** Precursor H&As were immunolabeled with Ptf1a and Tfap2a in E14.5 (left panels) and E17.5 (right panels) control and dcKO retinas. **B.** Nascent PHs were labeled by Blimp1 and Crx expression in E14.5 (left panels) and E17.5 (right panels) control and dcKO retinas. **A’.** Quantification of H&As in control and dcKO mutants at E14.5 and E17.5. **B’.** Quantification of PHCs in control and dcKO mutants at E14.5 and E17.5. Error bars represent S.D. **** p<0.0001, *** p<0.001, ** p<0.01, * p<0.05, ns: not significant. Scale bar applies to all panels in **A** and **B**.

We next examined the formation of the photoreceptor cells (PHCs) in the dcKO by immunostaining two PHC markers Prdm1 (Blimp1) and Crx. Prdm1 is expressed in fate-committed precursors of the PHC lineage, and Crx is expressed in differentiated PHCs (mostly cones at E14.5 and E17.5) ^87^. Overall, we observed a trend of increase of cells positive for the two markers at both E14.5 and E17.5, but only the increase in Crx+ cells at E14.5 reached statistical significance with a 1.58-fold change (**Fig. 3B and B’**).

The above findings suggest that SoxC factors regulate and coordinate the differentiation of the multiple early retinal lineages. Whereas RGCs and H&As were reduced in the dcKO retina, PHCs seemed to be increased, implying that the SoxC factors promote the RGC and H&A lineages, but repress the PHC lineage.

### Failure of RGC genesis in *Sox4/Sox11/Atoh7* tKO retinas

If the SoxC factors and Atoh7 function independently to promote the RGC lineage, we asked what would happen if all three genes were deleted. Accordingly, we examined RGC genesis in the tKO retina by immunostaining with the RGC markers mentioned above. At E14.5, Pou4f2+ and Isl1+ RGCs were almost completely abolished with 90.8% and 86.8% percent reductions, respectively, in the tKO retina. There were very few precursor RGCs in the NBL and almost no RGCs on the inner (basal) side where the presumed GCL was located (**Fig. 4A and A’**). The severe loss of RGCs was further confirmed by immunostaining for Pou4f1 and Ebf1, demonstrating that 90.8% of Pou4f1+ and 95.9% of Ebf1+ cells were abolished (**Fig. 4A and A’**). At E17.5, the tKO retina demonstrated a loss of 96.5% of Pou4f2+ cells and 92.1% of Isl1+ cells, eliminating both RGC precursors in the NBL and differentiating RGCs in the GCL (**Fig. 4B and B’**). In the same manner, RGCs in the GCL as marked by Pou4f1 and Ebf1 showed 91.3% and 96.0% reductions, respectively, in tKO retina (**Fig. 4B and B’**).

**Figure 4.**
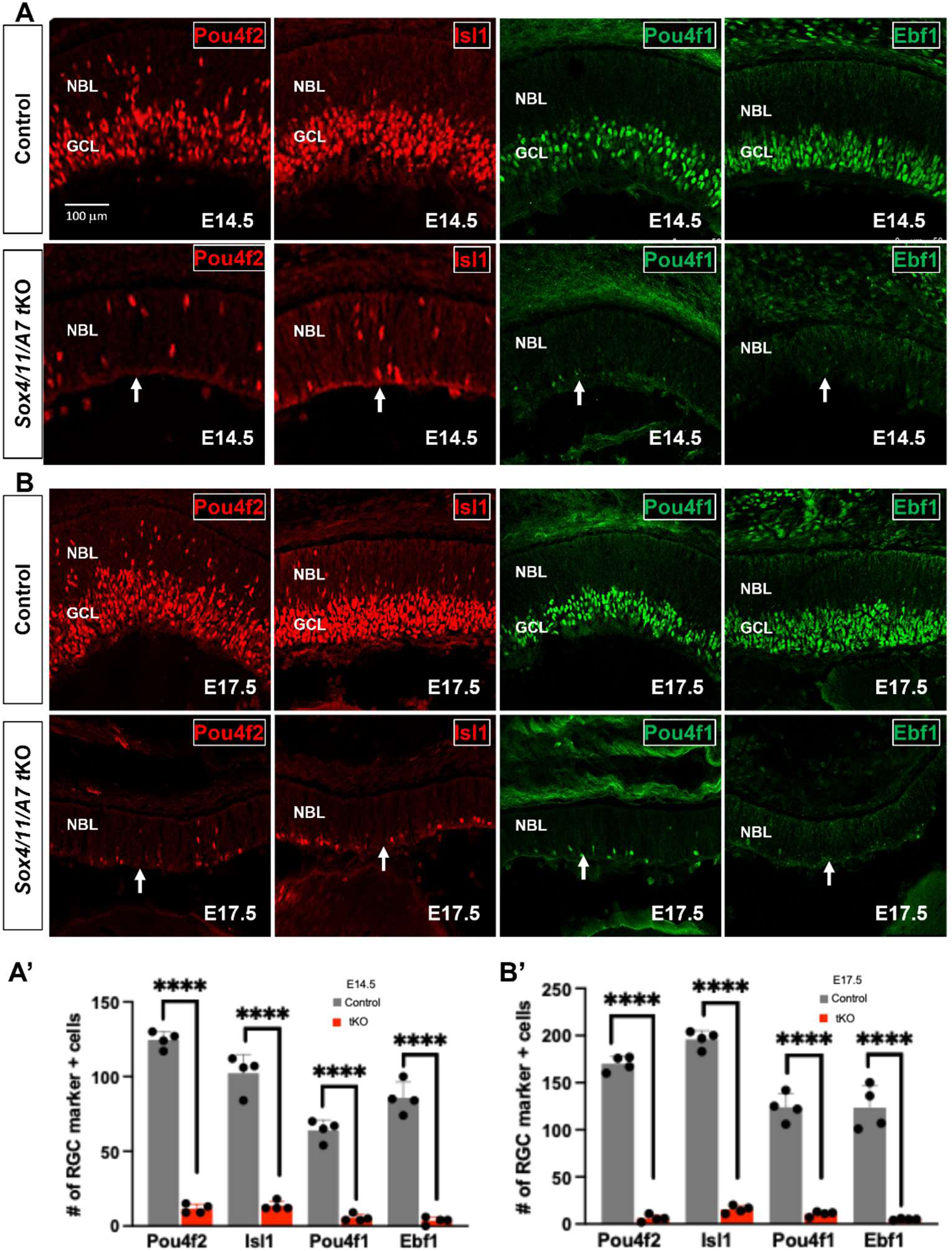
RGC lineage is abolished in the Sox4/Sox11/*Atoh7* tKO retina **A.** Immunofluorescence staining for four RGC markers in the E14.5 wild type (top) and tkO (bottom) retinas. Both precursor RGCs in the NBL as marked by Pou4f2 and Isl1, and RGCs in the GCL as marked by Pou4f1 and Ebf1 are almost completely depleted at E14.5. **B.** Labelling of RGCs for Pou4f2, Isl1, Pou4f1 and Ebf1 in E17.5 control and tKO retinas confirms complete loss of RGC precursors and more developed RGCs. **A’**. Quantification of RGC positive marker cells in control and tKOs at E14.5. **B’**. Quantification of RGC-specific markers at E17.5. Error bars represent S.D. **** p<0.0001. Scale bar applies to all panels in **A** and **B**. Arrows point to the missing GCL in the tKO retina.

The severe loss of RGC precursors was in stark contrast to the *Sox4/11* dcKO and *Atoh7*-null retinas, where precursor RGCs still form^3^ (**Fig. 2A, Suppl Fig. 5)**. The few remaining RGCs in the tKO retina could be due to residual *Sox4* and/or *Sox11* expression resulted from incomplete deletion by Cre, or the fact that *Sox12* still remained intact, but it was also possible that additional factors, such as Dlx1, Dlx2, and/or Pou3f1, were also involved in the RGC lineage formation ^15–17^. Nevertheless, our finding that the combined loss of Sox4, Sox11, and Atoh7 led to near complete elimination of both RGC precursors and differenting RGCs implies that the SoxC factors and Atoh7 constitute the major independent upstream regulatory inputs for establishing this lineage.

### Defect of other early retinal cell lineages in the *Sox4/Sox11/Atoh7* tKO retina

Similar to what was observed for the dcKO retina, horizontal and amacrine cell populations were also reduced in tKO retina. Ptf1a+ and Tfap2a+ cells were nearly absent in the E14.5 retina. On average, there was an 86% loss of Ptf1a+ cells and an 87.2% loss of Tfap2a+ cells (**Fig. 5A and A’**). At E17.5, however, the number of Ptf1a+ cells recovered, and no significant change was observed in the tKO retina as compared to the control (**Fig. 5A and A’**). Nevertheless, the innermost Tfap2a+ cells, which were amacrine cells, remained reduced by 53.1% and were localized in the inner retina due to the absence of the GCL. The outermost Tfap2a+ cells, which were horizontal cells, decreased by 75.4% in the tKO retina (**Fig 5A and A’**). The recovery of Ptf1a+ cells at E17.5 in both the dcKO and tKO retinas suggests a potential compensatory mechanism by other transcriptional regulators in the absence of Sox4/Sox11 and Atoh7.

**Figure 5.**
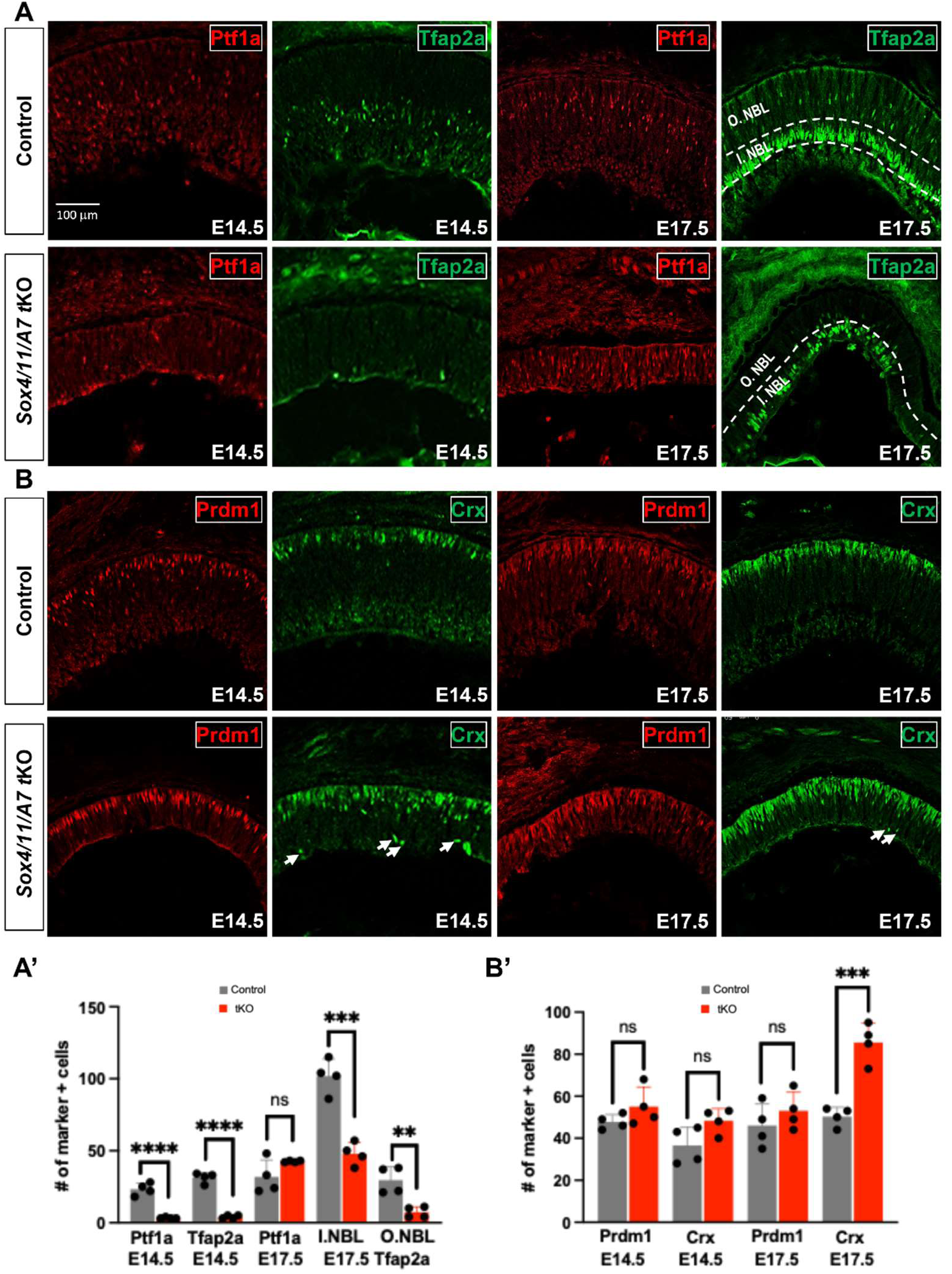
H&As are progressively reduced and PHs are upregulated in late tKOs **A.** Ptf1a and Tfap2a for H&A precursors in control and tKO retinas at E14.5 (left panels) and E17.5 (right panels). Dashed lines in the E17.5 Tfap2a panels demarcate the inner and outer NBL where horizontal and amacrine cells are located, respectively. **B.** Staining of Blimp1 and Crx for PHCs in control and tKOs at E14.5 (left panels) and E17.5 (right panels). White arrows point to Crx cells in tKOs at E14.5 and E17.5 that are ectopically located on the inner side of retina. **A’**. Quantification of Ptf1a and Tfap2a positive cells in control and tKOs at E14.5 and E17.5. **B’**. Quantification of Blimp1 and Crx positive cells at E14.5 and E17.5. Error bars represent S.D. **** p<0.0001, *** p<0.001, ** p<0.01, ns: not significant. Scale bar applies to all panels in **A** and **B**.

For the photoreceptor lineage, Prdm1+ and Crx+ PHCs showed a consistently upregulated trend in the tKO retina, again similar to what was observed in the dcKO retina. At E14.5, there was a 15.2% increase of Blimp1+ cells and a 32.2% increase of Crx+ cells, although the p values for both markers did not reach the 0.05 significance threshold. This upregulation trend persisted at E17.5, with Blimp1+ cells modestly elevated by 15.0%, but in contrast, Crx+ cells significantly increased by 70.1% (**Fig. 5B and B’**). Notably, some PHCs marked by Crx+ were ectopically located on the basal side in both the E14.5 and E17.5 tKO retinas, which were rarely seen in the dcKO retina (**Fig. 3B, 5B**). These findings suggest that when both the SoxC and Atoh7 inputs were absent, some RPCs might have been diverted to the photoreceptor fate.

### Decreased proliferation and increased cell death in the dcKO and tKO retinas

As noted above, both the dcKO and tKO retinas were substantially thinner than the controls. Also, RGC precursor cells were generated but lost in the dcKO retina. To investigate the underlying causes, we questioned whether cell proliferation and/or survival were compromised. We performed immunostaining for Ki67 and phosphorylated histone 3 (PH3), which are expressed in dividing RPCs; Ki67 marks all RPCs, whereas PH3 labels those in the M phase. We also performed EdU chase labeling to examine RPCs in the S phase. In the E14.5 dcKO retina, quantification of Ki67+, PH3+, and EdU+ cells exhibited 40.6%, 44.9%, and a 57.9% reductions respetively **(Fig.6 A and A’**). In the E17.5 dcKO retina, Ki67+ cells decreased by 21.8%, PH3+ cells were reduced by 45.3%, and EdU+ cells dropped by 66.0% (**Fig. 6A and A’**). We also performed TUNEL assay to evaluate whether cell survival was affected. In the dcKO retina, TUNEL+ cells had a 5.1-fold increase at E14.5, and a 5.0-fold increase at E17.5. Most of the TUNEL+ cells were localized to the innermost side of the retina, where the GCL would be (**Fig. 6A**), suggesting that the dying cells were likely RGCs and H&As. This was consistent with findings that these cell types were reduced in the dcKO retina.

**Figure 6.**
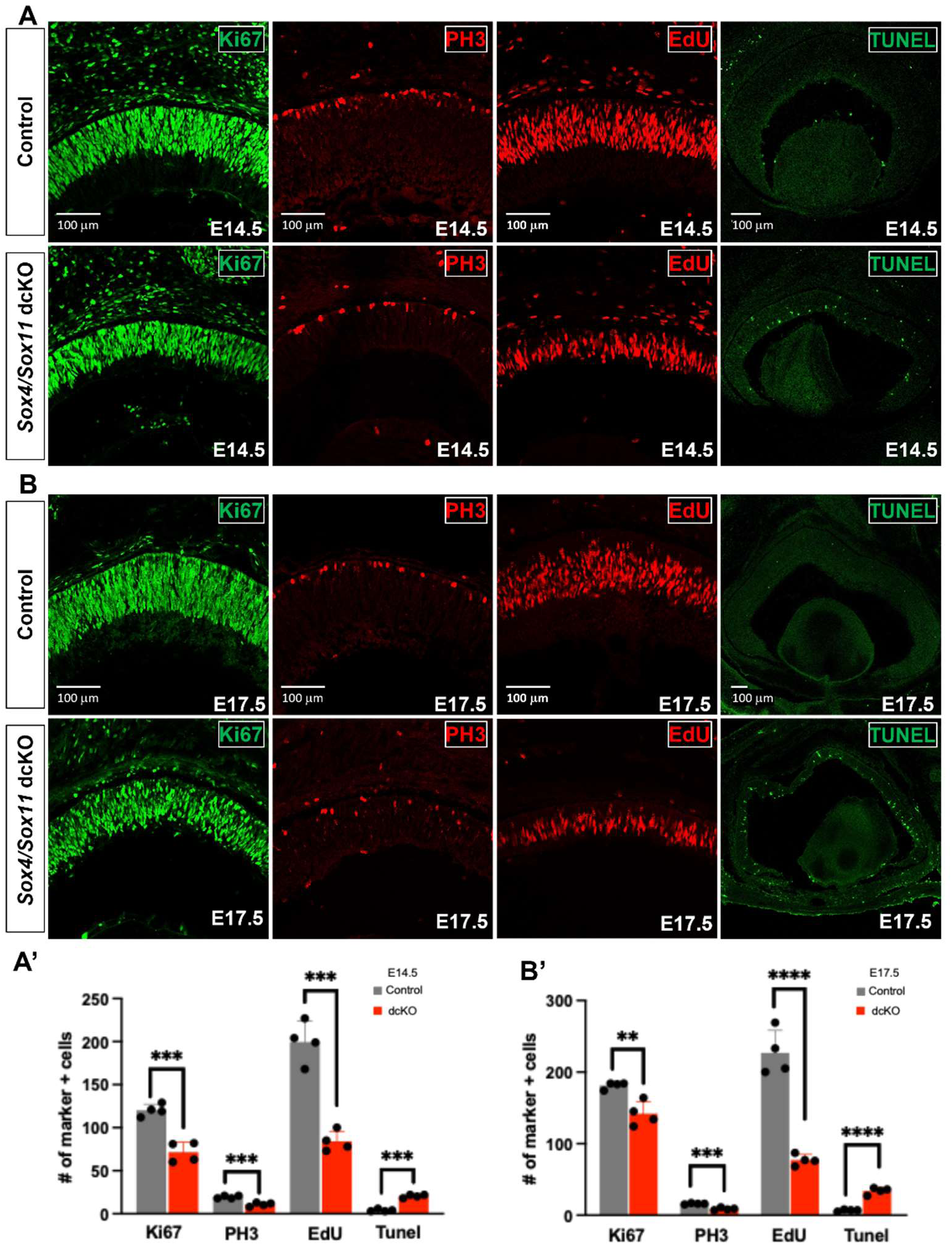
Decreased proliferation and increased apoptosis in the *Sox4*/*Sox11* dcKO retina A. Proliferating cells in the E14.5 control and dcKO retinas labeled by Ki67, expressed in dividing cells in all cell cycle phases, PH3, a mitotic cell marker, and EdU, incorporated during S phase. TUNEL assay detects apoptotic cells in control and dcKO retinas. **B.** Proliferating cells and apoptotic cells are labels in E17.5 control and dcKO retinas. **A’.** Quantification of Ki67, PH3, EdU and TUNEL positive cells in control and dcKO mutants at E14.5. **B’.** Quantification of Ki67, PH3, EdU, and TUNEL positive cells in control and dcKO mutants at E17.5. Error bars represent S.D.**** p<0.0001, *** p<0.001, ** p<0.01. Scale bars apply to corresponding pairs of control and mutant experiments.

Similar defects in proliferation and apoptosis were also observed in the tKO retina. Ki67+ cells, PH3+ cells, and EdU+ cells decreased by 17.0%, 40.0%, and 46.9%, respectively, at E14.5 (**Fig. 7A and A’**). The reduced proliferation continued at E17.5, with Ki67+ cells, PH3+ cells, and EdU+ cells reduced by 39.2%, 50.8%, and 52.0%, respectively (**Fig. 7B and B’**). Apoptotic cells in tKO retina increased by 3.4-fold at E14.5, and by 2.8-fold at E17.5. (**Fig. 7A and A’, Fig. 7B and B’**). In contrast to the dcKO retina (**Fig.** 6) or the *Atoh7*-null retina^13^ where dying cells are mostly located in the inner side, cell death in the tKO retina occurred throughout the full thickness, indicating cells died at earlier phases of differentiation, i.e., some nRPCs or tRPCs were likely dying. This difference also reflected our findings that most RGC precursors were absent in the tKO retina. One possible cause for the reduction in RPC proliferation is that secreted molecules from RGCs, such as Shh, Vegf, and Gdf11, which promote RPC proliferation and inhibit differentiation^80,81,88^ were lost due to the loss of RGCs in the dcKO and tKO. The SoxC factors may also directly regulate RPC proliferation and survival, as they are also expressed in proliferating RPCs ^3^ (**Fig. 1A**). Reduced proliferation and increased cell death likely both contributed to the thinning of the dcKO and tKO retinas.

**Figure 7.**
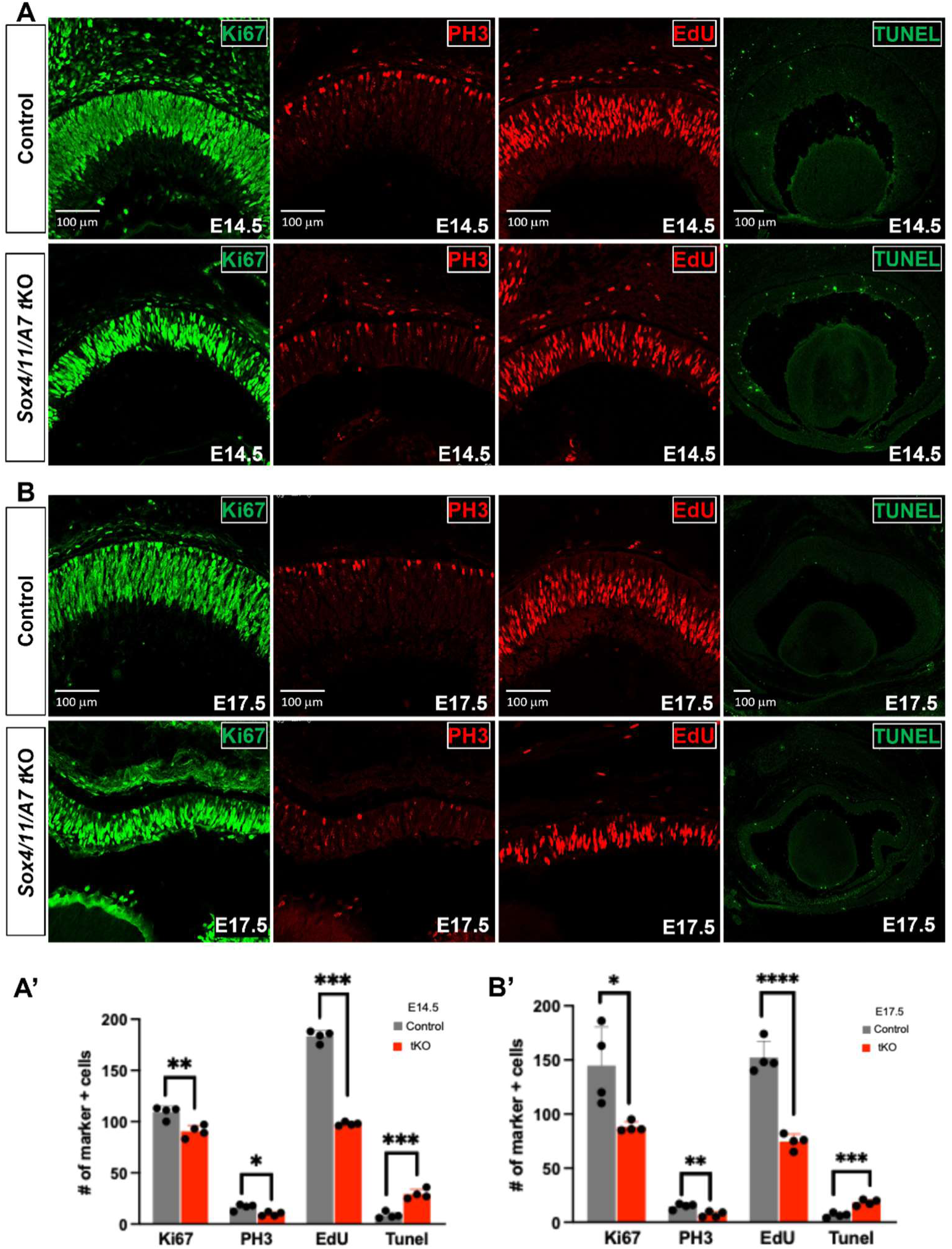
Decreased proliferation and increased apoptosis in the *Sox4*/*Sox11/Atoh7* tKO retina **A.** Labeling of proliferating cells by Ki67, PH3, and EdU, and apoptotic cells labeled by TUNEL in control and tKO retinas at E14.5. **B.** Labeling of proliferating cells by Ki67, PH3, and EdU, and apoptotic cells labeled by TUNEL in control and tKO retinas at E17.5. **A’.** Quantification of Ki67, PH3, EdU and TUNEL positive cells at E14.5. **B’.** Quantification of Ki67, PH3, EdU, and TUNEL positive cells at E17.5. Error bars represent S.D. *** p<0.001, ** p<0.01, * p<0.05. Scale bars apply to corresponding pairs of control and mutant experiments.

### Global transcriptomic changes in the dcKO and tKO retinas

To obtain a global view of downstream events and better understand how the SoxC factors interact with Atoh7 to regulate retinal cell differentiation, we performed bulk RNA-seq to identify transcriptomic changes in the dcKO and tKO retinas at E14.5 and E17.5. Differentially expressed genes (DEGs) between mutant and wild-type retinas were identified using the Wald test implemented in DESeq2 ^69^. Genes with an adjusted p-value (FDR) of 0.05 and an absolute log2 fold-change of 0.58 were considered differentially expressed.

In the E14.5 dcKO retina, we identified 1,797 downregulated and 2,089 upregulated genes (**Fig. 8A, Suppl Table 1**). The downregulated DEGs included core RGC transcription factor genes, such as *Pou4f2, Isl1, Pou4f1, Ebf1, Ebf3, Dlx1, Dlx2* and *Pou3f1,* as well as genes encoding proteins for RGC structure and function, including *Sncg, Stmn2, Gap43, Nefl, Nefm,* and *Ina* ^3,44,45,89–91^. This was consistent with the defective RGC development in the dcKO retina as revealed by marker analysis. H&A marker genes such as *Ptf1a*, *Tfap2b*, *Gad2*, *Slc32a1*, and *Prdm13* ^3^were also drastically downregulated (**Fig. 8A, Suppl Table 1**), whereas many genes encoding proteins for photoreceptor (cone) differentiation and function (e.g. *Crx*, *Gnat2*, *Thrb*, *Klc3*, *Gnb3*, *Pdc*, *Rbp3*, *Arr3, Gngt2, Cngb3,* and *Atp1b2*) ^3,92,93^ were significantly upregulated, which was again consistent with the changes of the two lineages observed from the IHC analysis. In the E17.5 dcKO retina, there were a total of 2,481 downregulated and 2,440 upregulated DEGs (**Suppl Table 2**). Transcriptional changes largely mirrored those observed at E14.5, with sustained downregulation of RGC and H&A programs, and persistent upregulation of photoreceptor genes (**Fig. 8A**). Many RGC genes such *Nefl* and *Nefm* showed stronger downregulation compared to the E14.5 dcKO retina, reflecting the continued loss of RGCs (**Fig. 8A**). H&A marker genes such as *Tfap2a/b*, *Lhx1*, *Gad1, Slc32a1*, *Onecut1/3, Barhl2* and *Nr4a2* continued to be downregulated (**Fig. 8A, Suppl Table 2**). Interestingly, both *Atoh7* and *Ptf1a* were upregulated (**Fig. 8A, Suppl Table 2**), likely due to a negative feedback mechanism^80,81^. The discrepancy in the changes of Ptf1a with the IHC (**Fig. 3A**) results may have resulted from the different sensitivities of the assays, but the RNA-seq data should be more reliable. Cone genes such as *Crx*, *Neurod4*, *Gnat2*, *Thrb*, *Klc3*, *Gnb3*, *Pdc*, *Rbp3*, *Arr3, Gngt2, Cngb3,* and *Atp1b2* remained to be upregulated (**Fig. 8A, Suppl Table 2**). Additionally, many rod genes such as *Nr2e3, Nrl* and *Rho* were upregulated in the E17.5 dcKO retina (**Suppl Table 2**). The upregulation of rod genes suggests that precocious rod development occurred in the dcKOs since these genes, particularly *Rho*, are normally expressed at later stages ^94^. These results confirmed that the SoxC factors coordinate the production of the early retinal cell types. The findings also support a previous report indicating the SoxC factors regulate the different timings of cone versus rod production ^59^.

**Figure 8.**
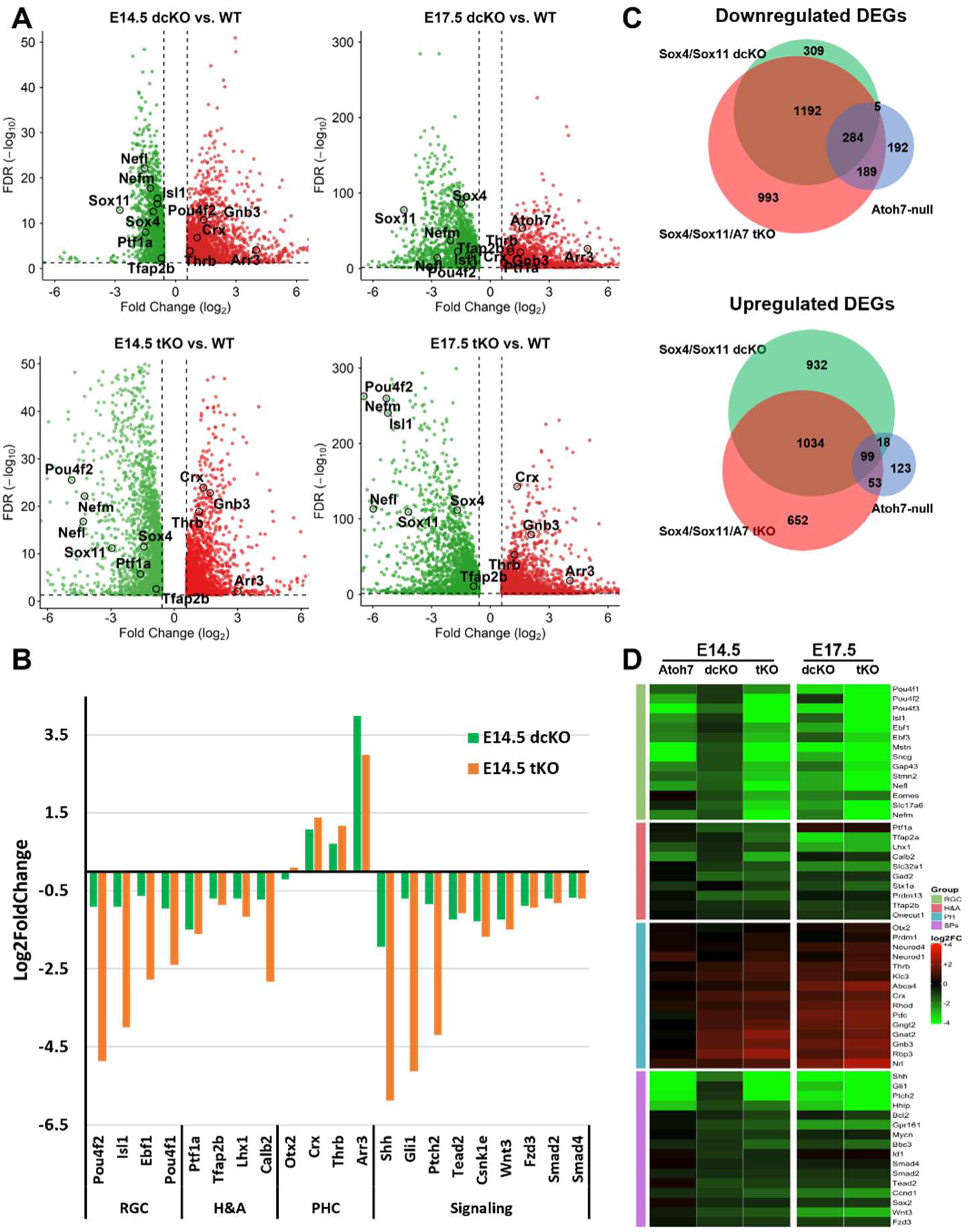
Global gene expression changes in dcKO and tKO retinas reveal impact on retinal lineages and developmental pathways **A.** Volcano plots of downregulated (green) and upregulated (red) DEGs in E14.5 and E17.5 dcKO and tkO retinas. Select lineage specific genes are highlighted. **B.** Bar graph displaying different degree of changes of select lineage specific marker genes and genes of signaling pathways in the E14.5 dcKO and tKO retinas. **C.** Venn diagrams displaying the overlaps of the DEGs in the three mutant retinas, dcKO, *Atoh7*-null, and tKO. Downregulated and upregulated DEGs are separately compared. **D.** Heatmap depicting changes of select lineage and signaling pathway genes in *Atoh7*-null, *Sox4/Sox11* dcKO, and *Sox4/Sox11/Atoh7* tKO retinas at E14.5 and E17.5. E17.5 Atoh7-null data is not available.

In the E14.5 tKO retina, a total of 2,663 genes were downregulated and 1,849 upregulated relative to controls (**Fig. 8A**). Changes of lineage-specific genes were consistent with the IHC marker analysis, with RGCs and H&As downregulated and photoreceptor genes upregulated (**Fig. 8A, Suppl. Table 3**). At E17.5, 2,808 downregulated DEGs and 2,621 upregulated DEGs were identified (**Fig. 8A, Suppl Table 4**). These DEGs continued to reflect the changes in the three lineages, decreases in the RGC and H&A lineages, and expansion and precocious differentiation of the photoreceptor lineage (**Fig. 8A, Suppl Table 4**). Notably, essentially all the RGC genes, e.g., *Pou4f2, Isl1, Pou4f1, Ebf1, Ebf3, Dlx1, Pou3f1 Sncg, Stmn2, Gap43, Nefl, Nefm,* and *Ina*, exhibited substantially greater reduction in the tKO retina than in the dcKO retina (**Suppl. Tables 1-4)**, which was evident in their fold change distributions along the x axis of the volcano plots for both E14.5 and E17.5 retinas (**Fig. 8A**). Genes involved in the H&A lineage, such as *Ptf1a*, *Lhx1*, *Tfap2b,* and *Calb2,* were also reduced in the E14.5 tKO retina, but only to slightly greater degrees as compared to the dcKO retina. PHC genes such as *Crx*, *Neurod1*, *Neurod4*, *Gnb3*, and *Thrb* displayed substantial increases in expression, and the increases were largely comparable to those observed in the dcKO retina (**Fig. 8A, Suppl. Tables 1-4**). The quantitative change differences in the dcKO and tKO retinas were better illustrated by the bar charts of selected lineage specific DEGs (**Fig. 8B, Suppl Fig. 6**), confirming the changes of the three lineages, particularly the different degrees of RGC loss, in the dcKO and tKO retinas.

### Shared and distinctive roles of SoxC factors and Atoh7 in regulating retinal cell differentiation

The differences in cell differentiation and gene expression between the dcKO and tKO retinas indicated that the SoxC factors and Atoh7 played differential roles in the three early retinal lineages. To further understand the interaction of the SoxC factors and Atoh7, we compared the E14.5 dcKO and tKO DEG lists with the *Atoh7*-null DEG list we previously reported ^3^. These comparisons revealed overlapping and distinct sets of DEGs in the different mutant retinas. As illustrated by the Venn diagrams (**Fig. 8C**), there were extensive overlaps between the three mutant retinas within the downregulated gene sets. A core group of 284 genes was commonly reduced across all three genotypes, including those encoding canonical RGC specification and differentiation regulators as well as structural and functional proteins such as *Pou4f2, Isl1*, *Pou4f1*, *Ebf1*, *Dlx1*, *Pou3f1*, *Sncg*, *Nefl*, *Nefm*, *Gap43, Ina*, and *Stmn2*. This shared loss in gene expression underscores that the SoxC factors and Atoh7 converge on a transcriptional program essential for establishing and maintaining the RGC identity. The differences in RGC gene changes between the dcKO and tKO retinas at E14.5 and E17.5 could be further illustrated by the heatmap in **Fig. 8D**. Similarly, many of these RGC genes also showed much larger changes in the tKO retina than in the *Atoh7*-null retina (**Fig. 8D**). This was consistent with the findings that Sox4/Sox11 or Atoh7 alone still permitted limited formation of RGC precursors, but the gene expression program was compromised. In contrast, combined loss of the SoxC factors and Atoh7 causes a close to complete failure of RGC genesis, and thus substantially larger decrease in the expression of the RGC genes (**Fig. 8D**).

Together, these data support the cooperative roles for Sox4/Sox11 and Atoh7 as two major and maybe only upstream regulatory inputs in driving tRPCs to the RGC fate by activating the RGC gene expression program. Notably, individual genes were differentially affected in the dcKO and *Atoh7*-null retinas, indicating each gene received different levels of inputs from the two regulatory pathways. The H&A genes also demonstrated differential changes in the three genotypes. At E14.5, only a few of them demonstrated downregulation in *Atoh7*-null retina, whereas many showed significant downregulation in the *Sox4/Sox11* dcKO retina. The changes in the E14.5 tKO retina seemed to be the sum of changes in the *Atoh7*-null and dcKO retinas (**Fig. 8D**). At E17.5, the dcKO and tKO retinas displayed very similar changes in these genes (**Fig. 8D**). These results indicated that, whereas both participated in the establishment of the H&A lineage, the SoxC factors likely played a much larger role than Atoh7. As mentioned above, *Ptf1a* was substantially downregulated at E14.5 in both dcKO and tKO retinas (**Fig. 8D**), but this change diminished at E17.5, demonstrating the complexity of the process and indicating that other upstream factors such as Foxn4^24^ may compensate for the loss of the SoxC factors. For the PHC lineage, there was upregulation of multiple PHC genes in all three mutant retinas, with *Atoh7*-null retina demonstrating the smallest, and the tKO retina showing the largest changes (**Fig 8D**), indicating that the SoxC factors also played a larger role than Atoh7 in repressing this lineage. For the upregulated DEGs, the overlaps between the *Atoh7*-null list and those of the other two mutants were very small (**Fig 8C**), indicating Atoh7 and the SoxC factors repress different molecular pathways, either directly or indirectly, which emphasized the differences of the SoxC factors and Atoh7 in regulating retinal development.

We also performed gene ontology (GO) analysis of the dcKO and tKO DEGs to gain further insights into the downstream events of Sox4/11 and Atoh7. For simplicity, we combined E14.5 and E17.5 DEGs for each genotype. As might have been expected, the top 20 enriched biological process GO terms for the downregulated DEGs in both the dcKO and tKO retinas were similar and all related to neural development, with the top enriched terms including “generation of neurons” and “neuron differentiation”. KEGG pathway analysis further confirmed this with “axon guidance” being one of the top enriched pathways (**Suppl Fig. 7**). Notably, “cell cycle” and “hippo signaling pathway” were also among the top pathways whose components were significantly enriched in the downregulated DEG lists. This was consistent with the reduced RPC proliferation observed in both the dcKO and tKO retinas. Additionally, component genes of the Shh signaling pathway ^3^, which modulate RPC proliferation via a feedback mechanism, and *Sox2*, which directly regulates RPC proliferation ^38^, were also downregulated in the dcKO and tKO retinas (**Suppl Tables 1-4**). These findings emphasized that the SoxC factors participate in both the intrinsic and extrinsic mechanisms coordinating proliferation and differentiation in the developing retina. GO terms in the upregulated gene lists further confirmed the upregulation of the PHC lineage, as reflected by enriched biological process GO terms such as “visual perception” and KEGG pathways such as “phototransduction”, “calcium signaling pathway”, and “glutamatergic synapse”. There were additional enriched pathways, including “ECM-receptor interaction”, “Wnt signaling pathway”, “PI3K-AKT signaling pathway”, and “MAPK signaling pathway” (**Suppl. Fig. 7**). Although the biological significance of these pathways required further investigation, they demonstrate that the SoxC factors, either alone or together with Atoh7, regulate retinal cell differentiation via multiple molecular processes.

### SoxC factors directly regulate genes expressed in multiple cell states and lineages

The DEGs identified likely included both direct and indirect targets of the SoxC factors. To identify the direct target genes, we attempted to perform CUT&Tag ^36,77^ to identify the binding sites for Sox4 and Sox11 in the E14.5 retina, but were only able to obtain reliable results for Sox11. Nevertheless, given the high similarity of the DNA-binding domains and the functional redundancy of the two factors, the two transcription factors likely share similar if not identical targets. We also performed CUT&Tag for the active enhancer mark H3K27ac ^95^ to identify the active enhancers in the E14.5 retina. Using MACS2, we identified a total of 39,145 Sox11 peaks (**Suppl Table 5**) and 64,838 H3K27ac peaks (**Suppl Table 6**). As we did previously, we intersected the Sox11 peaks with the peaks identified by scATAC-seq and narrowed down the active enhancers bound by Sox11 to 12854 and identified their cell state/type specificities^36^ (**Suppl. Table 5**). The activities of these Sox11-bound enhancers as defined by the cutting frequencies in scATAC-seq in the different cell states/types did not demonstrate high cell state/type specificities (**Fig. 9A**). This was in contrast to other transcription factors such as Atoh7, Otx2, Pou4f2 and Isl1, which bind enhancers that have strong cell state/type specificity ^36^. Nevertheless, this was consistent with the RNA-seq results showing that genes expressed in multiple cell states/types were affected in the mutant retinas. Motif enrichment analysis by HOMER of regions ±250 bp of the called summits indicated that the top enriched motif was similar to previously reported the Sox4/Sox11 binding motifs ^52^ (**Fig. 9B**), further supporting the validity of the experiment. 23.9% of the Sox11 bound regions contained the consensus site (log p value of enrichment = −1.025e+03), which was similar to previously reported ChIP-seq results for the SoxC factors in other tissues ^57,96–98^.

**Figure 9.**
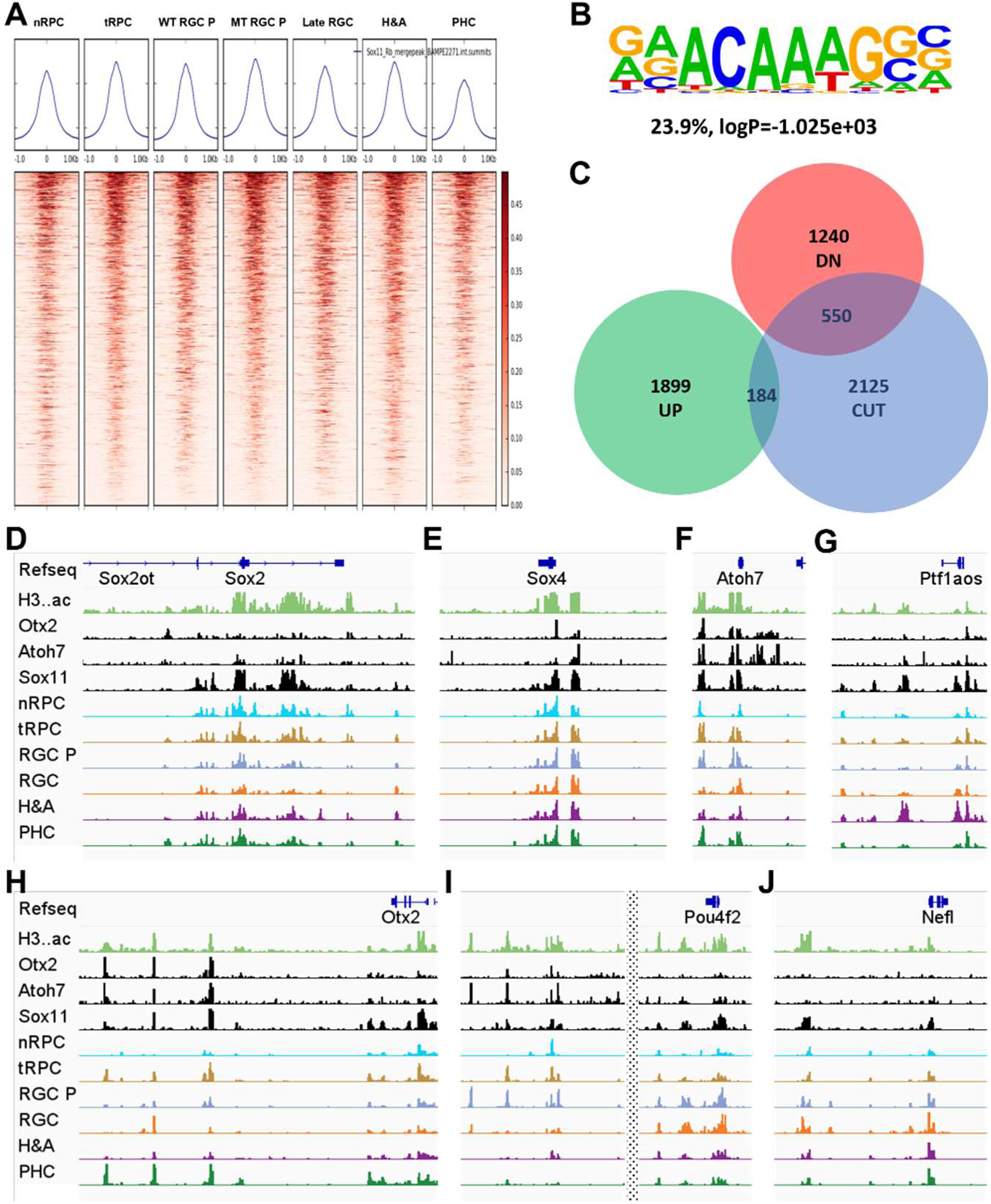
CUT&Tag identify genomic regions bound by Sox11 and its target genes **A.** Intersection of CUT&Tag and scATAC-seq peaks demonstrate the cell type/state specificities of Sox11 bound regions. **B.** Logo of the top enriched motif from the Sox11 bound regions resemble previously reported Sox4/Sox11 bound motif. **C.** Overlap of Sox11 target genes with DEGs in the E14.5 dcKO retina. **D-J.** Genome tracks of CUT&Tag peaks for H3K27ac (H3..ac), Otx2, Atoh7, and Sox11, and scATAC-seq peaks reported previously for key regulatory and cell type-specific genes. The scATAC-seq peaks demonstrate the cell type/specific opening of the associated regulatory elements. RGC P=RGC precursors.

We then identified the genes associated with these active Sox11-bound enhancer regions based on the peak-to-gene links we previously published ^36^. A total of 2859 plausible direct target genes were identified (**Suppl Table 5**). Top enriched GO terms and pathways of these target genes included “nervous system development”, “neurogenesis”, “hippo signaling pathway”, “axon guidance”, and “cell cycle” (**Suppl Fig. 8**), consistent with enriched GO terms of DEGs in the dcKO and tKO retinas. Further, the identified target genes overlapped with the DEGs in the E14.5 dcKO retina, including 550 down regulated genes and 184 up regulated genes (**Fig. 9C**). Importantly, the target genes included those encoding key regulatory factors functioning in the different cell states and types at this developmental stage, such as Ccnd1, Sox2, and Tead1 in proliferating nRPCs, Atoh7, Otx2, Sox4, Sox11, and Neurod1 in transitional RPCs, Isl1, Pou4f2, Ebf1, Pou3f1, and Klf7 in RGCs, Crx and Neurod4 in photoreceptors (cones), and Ptf1a, Tfap2a, Tfap2b, Prox1, Bhlhe22, and Lhx1 in H&As (**Suppl Table 5**). These findings strongly suggested that these genes were bona fide targets and that the SoxC factors directly regulated diverse processes in the developing retina.

Comparison of the Sox11 bound enhancers with the H3K27ac enriched regions and the enhancers bound by Atoh7 and Otx2 from our previous study revealed several insights (**Fig. 9D-J**). First, essentially all the regions bound by Sox11, Atoh7, and Otx2 overlapped with the H3K27ac regions, indicating these sites are all active enhancers (**Fig. 9D-J**). Second, the Sox11 bound enhancers associated with nRPCs and tRPCs genes often remained open in the differentiated cells such as RGCs, when the genes are turned off (see enhancers associated with *Sox2* and *Atoh7*) (**Fig. 9D, H**). Third, Sox11-bound enhancers associated with nRPC genes (e.g. *Sox2*) tend not to be bound by Atoh7 or Otx2 (**Fig. 9D**). This was consistent with the fact that Atoh7 and Otx2 are not expressed in nRPCs. Fourth, genes involved in specific lineage, such as *Sox4*, *Atoh7* and *Pou4f2* for RGCs (**Fig. 9E, F, I**), *Ptf1*a for H&As (**Fig. 9G**), and Otx2 (**Fig. 9H**) and *Crx* (not shown) for PHCs, were often regulated by Sox11, Atoh7, and Otx2 together; this was achieved by either co-binding the same enhancers or binding to distinct enhancers. This further supported the notion that transcription factors expressed in the tRPCs function together, either collaboratively or competitively, to achieve the eventual outcome of lineage specific gene expression. Fifth, whereas Sox11 and Atoh7 co-regulated the RGC precursor genes such as *Pou4f2* (**Fig. 9I**) and *Isl1* (not shown), Sox11 continued to regulate genes expressed in differentiating RGCs, such as *Nefl* (**Fig. 9J**), when *Atoh7* is inactivated. These findings indicated that Sox11, and likely Sox4 as well, directly regulate gene expression in multiple cell states/types and further confirmed the broad cellular functions of the SoxC factors in the developing retina.

## Discussion

In this study, we thoroughly investigated the roles of the SoxC factors (Sox4 and Sox11) in early retinal development. Our major focus is on the positions of these factors in the gene regulatory network for RGC genesis and their relationships to other key transcription factors, particularly Atoh7. We not only validated the critical roles of the SoxC factors, Sox4 and Sox11, in promoting RGC genesis, but also established that they function both independently of, and in cooperation with, Atoh7 in the process. Previous studies suggest that factors other than Atoh7 participate in establishing the RGC lineage from transitional RPCs, and our current study reveals that the SoxC factors fulfill that role. The SoxC factors and Atoh7 are both critically required for RGC genesis but are largely independent of each other for expression. There is upregulation of Atoh7 in the dcKO retina, but that likely occurs via the feedback mechanism ^80–82^. The SoxC factors and Atoh7 function within tRPCs, where they promote early neurogenic competence and activate the gene expression program for the RGC lineage. Consistent with this framework, loss of either the SoxC factors or Atoh7 results in a marked reduction of RGCs, but RGC precursors still form. Nevertheless, when both the SoxC factors and Atoh7 are absent, the RGC lineage fails to form. These findings indicate that the SoxC factors and Atoh7 represent two major, if not the only, upstream regulatory inputs required for establishing the RGC lineage (**Fig. 10**). In line with this, we observed overlapping sets of downregulated RGC associated genes across *Atoh7*-null, *Sox4/Sox11* dcKO, and *Sox4/Sox11/Atoh7* tKO retinas, including genes linked to RGC differentiation and axon growth, with the tKO retina demonstrating the largest decrease in the RGC gene expression program. These results indicate that both inputs are needed to fully activate the RGC gene expression program. Genome-wide DNA-binding mapping of Sox11 further confirmed that the SoxC factors and Atoh7 function together to fully activate the early regulatory RGC genes, including *Pou4f2*, *Isl1*, *Dlx1*, *Dlx2* and *Pou3f1*, so that the RGC differentiation program can be activated. On the other hand, differential changes of individual genes in the dcKO and *Atoh7*-null retinas suggest the dependence of each gene on the SoxC factors and Atoh7 is different. Of note, whereas Atoh7 expression is transient and is not maintained in differentiated RGCs, the SoxC factors persist, suggesting that whereas both SoxC and Atoh7 promote RPC competence and entry into the RGC trajectory, the SoxC factors likely provide continued transcriptional support needed for further differentiation and maturation.

**Figure 10.**
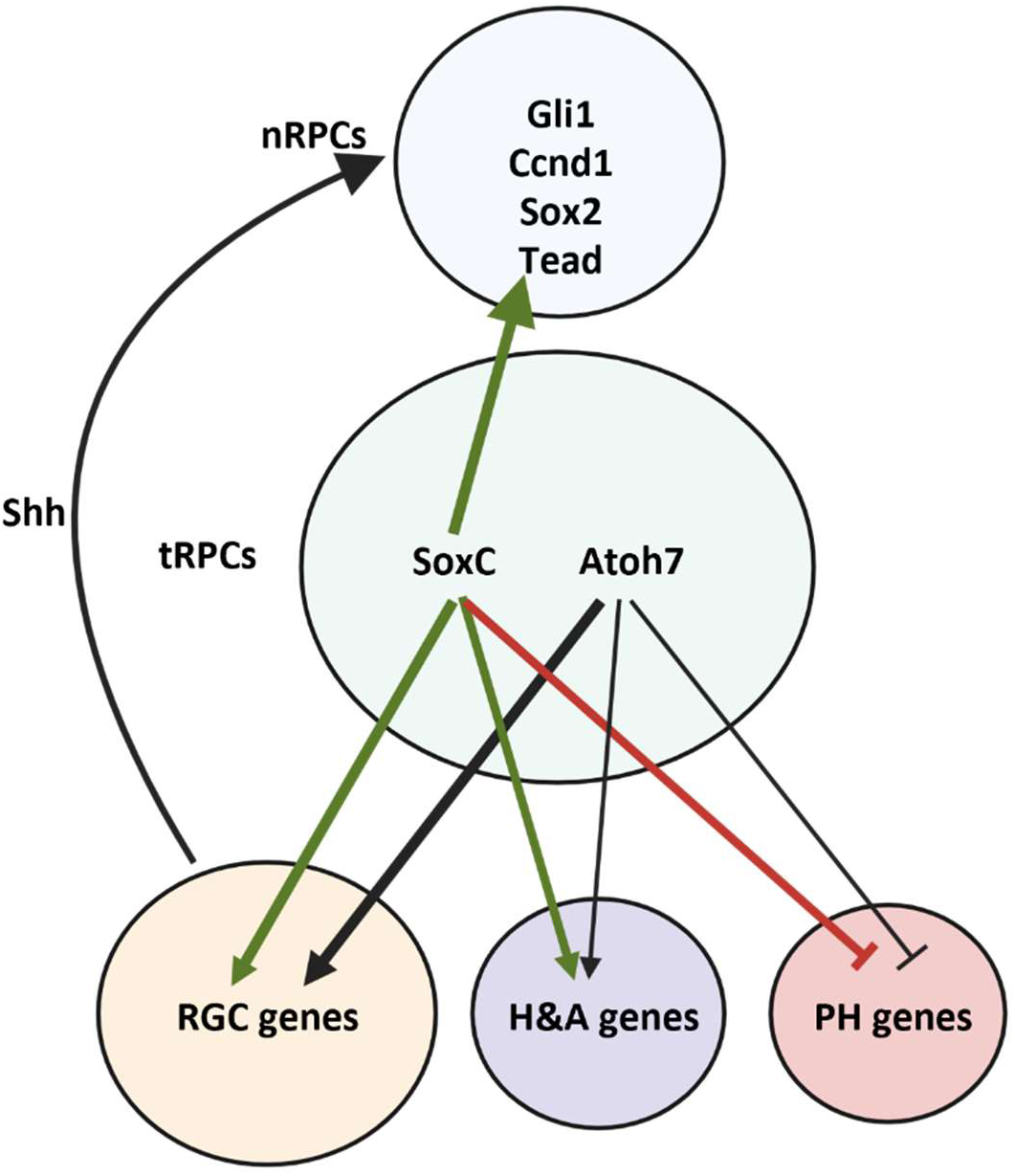
A model of the SoxC and Atoh7 regulatory programs during early retinal development. SoxC (Sox4 and Sox11) and Atoh7 act within transitional retinal progenitor cells (tRPCs) to shape early fate specification. Green and black arrows (arrowhead) indicate activation, showing that SoxC and Atoh7 promote expression of RGC genes and contribute to activation of horizontal/amacrine (H&A) genes. Red lines with a horizontal bar indicate repression, illustrating SoxC mediated suppression of the photoreceptor (PH) gene program to prevent premature PH differentiation. SoxC dependent changes are also linked to progenitor state regulation, including effects on Hippo/Shh pathway genes in naïve RPCs (nRPCs), both directly and indirectly. Overall, the diagram summarizes SoxC and Atoh7 dependent transcriptional control of early retinal developmental programs.

Another important finding from our study is that, unlike Atoh7, the SoxC factors play major roles in the other early retinal lineages. In both the dcKO and tKO retinas, genes involved in H&A differentiation program are reduced at E14.5 (**Fig. 10**). Interestingly, H&A precursor cells expressing Ptf1a recovered at E17.5 in both the dcKO and tKO retinas, although the differentiated H&A cells remain reduced. This was confirmed by the RNA-seq analysis. This finding indicated that other regulatory inputs exist to compensate for the loss of the SoxC factors at later developmental times. In contrast, loss of Atoh7 resulted in just minor changes in H&As at both the cellular and transcriptomic levels, indicating it plays only minor roles in this lineage. For the photoreceptor (PHC) lineage, however, loss of Sox4 and Sox11 leads to increased cone and precocious rod differentiation, indicating that SoxC factors normally repress photoreceptor development. This was also validated by the transcriptomic analysis, showing that many photoreceptor genes in the dcKO and tKO retinas, including components of key pathways involved in photoreceptor function and vision signal processing, were increased. Particularly of note is that there is precocious expression of rod genes in the absence of the SoxC factors. These findings suggest that at the early stages of retinal development, the SoxC factors function to negatively modulate the overall photoreceptor program as well as to repress rod gene expression. In contrast, Atoh7 seems to play a relatively minor role in the PHC lineage (**Fig. 10**). Thus, the SoxC factors function to coordinate the production of the different retinal lineages.

Our data also point to a broader influence of SoxC on the progenitor cells. The observed defects in RPC proliferation and survival in the dcKO and tKO retinas may reflect both direct and indirect roles of the SoxC factors (**Fig. 10**). Some key regulator genes, such as Sox2 and Ccnd1, are directly regulated by the SoxC factors. The SoxC factors may also directly regulate nRPC proliferation by modulating the Hippo pathway. This again is different from Atoh7 since Atoh7 is not expressed in proliferating nRPCs and thus does not directly modulate their proliferation. On the other hand, both the SoxC factors and Atoh7 influence nRPC proliferation indirectly via a feedback mechanism. Similar to Atoh7, SoxC factors promote the expression of Shh in RGCs, which is released and then acts on nRPCs, leading to increased proliferation and reduced RGC production via downstream effectors such as Gli1 and Ccnd1 (cyclin D1) (**Fig.10**). This mechanism likely contributes to the reduced proliferation as well as the increased RGC precursors observed in the dcKO and tKO retinas. Other signaling pathways such as the Wnt pathway, the PI3K-AKT pathway, and the MAPK pathway may also be involved as indicated by their disruption in the mutant retinas, but how these pathways integrate into the overall regulatory mechanisms controlling retinal development requires further investigation. Notably, there is also increased RPC death in the mutant retinas, particularly in the tKO retina, but the mechanism leading to their death is not known. The dying RPCs are likely tRPCs normally destined to specific lineages such as RGCs and H&As. If that is the case, why do they die instead of adopting a different fate, e.g. PHCs, is an interesting and important question that warrants further investigation, as it is pertinent to the central issue of cell fate choices during retinal development. It is worth noting that although the current study focuses on early retinal development, the SoxC factors likely also regulate late cell lineages as indicated by their continued expression at late stages.

The different roles of the SoxC factors and Atoh7 in regulating the non-RGC cell states/types demonstrate the complex but interconnecting mechanisms underlying the coordinated generation of the different retinal cell types. Overall, our study reveals the broad roles of the SoxC factors in regulating retinal development at different levels in various cell states/types. The SoxC factors achieve their regulatory roles by regulating target genes in different cell states/types as indicated by our CUT&Tag analysis. We have now firmly established that SoxC factors collaborate with Atoh7 in establishing the RGC lineage, but they may collaborate with different partners in the other lineages, which remain to be identified. How the SoxC factors interact and influence the epigenetic landscape to carry out their broad functions remains to be clarified. Our finding that many of the Sox11 bound enhancers remain open in cell states in which the relevant gene is turned off is interesting. This is consistent with our previous observation that enhancers associated with many cell cycle genes tend to be in an open state in all cell states/types^36^. This discordance between enhancer openness and gene expression has been observed, suggesting that openness is not equivalent to activity ^99,100^. It is worth noting that a relatively low percentage of regions bound by Sox11 contain the consensus Sox4/Sox11 binding sites. This is unlikely to be caused by experimental errors, such as non-specific binding of the antibody, since these regions make biological sense and are associated with genes involved in retinal development. Also, this number is consistent with DNA-binding mapping data for Sox4 and Sox11 in other tissues ^57,96–98^. Given the various functions of the SoxC factors in retinal development, they may bind to diverse DNA sequences with different affinities via collaborating with distinct factor(s) in various cellular contexts. It is also possible that they interact with DNA indirectly via association with other transcription factors.

## Acknowledgments

We are grateful to Dr. Fuguo Wu for technical advice. We thank Dr. Elisabeth Sock for generously providing the Sox4 and Sox11 antibodies. We also thank Dr. Donald Yergeau at the Genomics and Bioinformatics Core of the University at Buffalo for sequencing the bulk RNA-seq and CUT&Tag libraries. Imaging and cell analysis were carried out in the Confocal Microscopy and Flow Cytometry Facility (CMFCF) at the University at Buffalo. We thank Dr. Wade J. Sigurdson for providing help and advice during imaging. We further acknowledge the Department of Laboratory Animal Resources (LARS) animal facility in the Molecular Research Complex (MRC) at Roswell Park Comprehensive Cancer Center for excellent animal care and maintenance of our mouse colonies. This work was funded by grants (R01EY020545, R01EY029705) from the National Eye Institute to XM. The content is solely the responsibility of the authors and does not necessarily represent the official views of the funding agencies.

## Author contribution statements

SE designed and performed the experiments, collected data, performed bioinformatic analysis of the bulk RNA-seq data, and co-wrote the paper; YG performed experiments and collected data; NN performed bioinformatic data of the CUT&Tag data; VL provided the Sox4 and Sox11 mouse lines and provided technical advice; TL guided the bioinformatics analysis; XM conceived the project, obtained funding, analyzed data, and co-wrote the paper.

## Supplementary Figure Legends

**Supplementary Figure 1.**
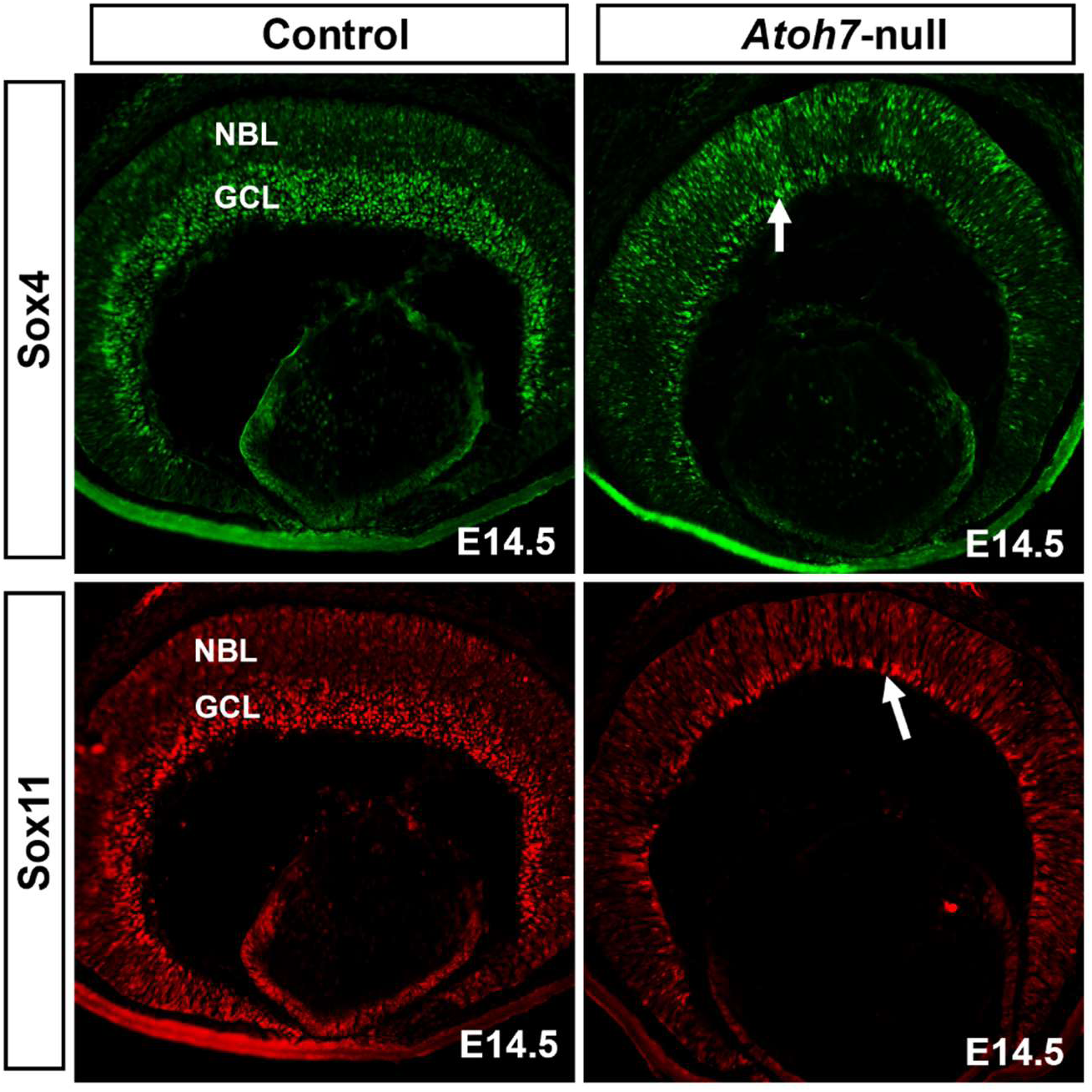
Immunofluorescence staining showing that Sox4 and Sox11 expression in the NBL is not affected in the E14.5 *Atoh7*-null retina. Note that the GCL, where Sox4 and Sox 11 are also expressed in the control retina, is largely gone in the *Atoh7*-null retina.

**Supplementary Figure 2.**
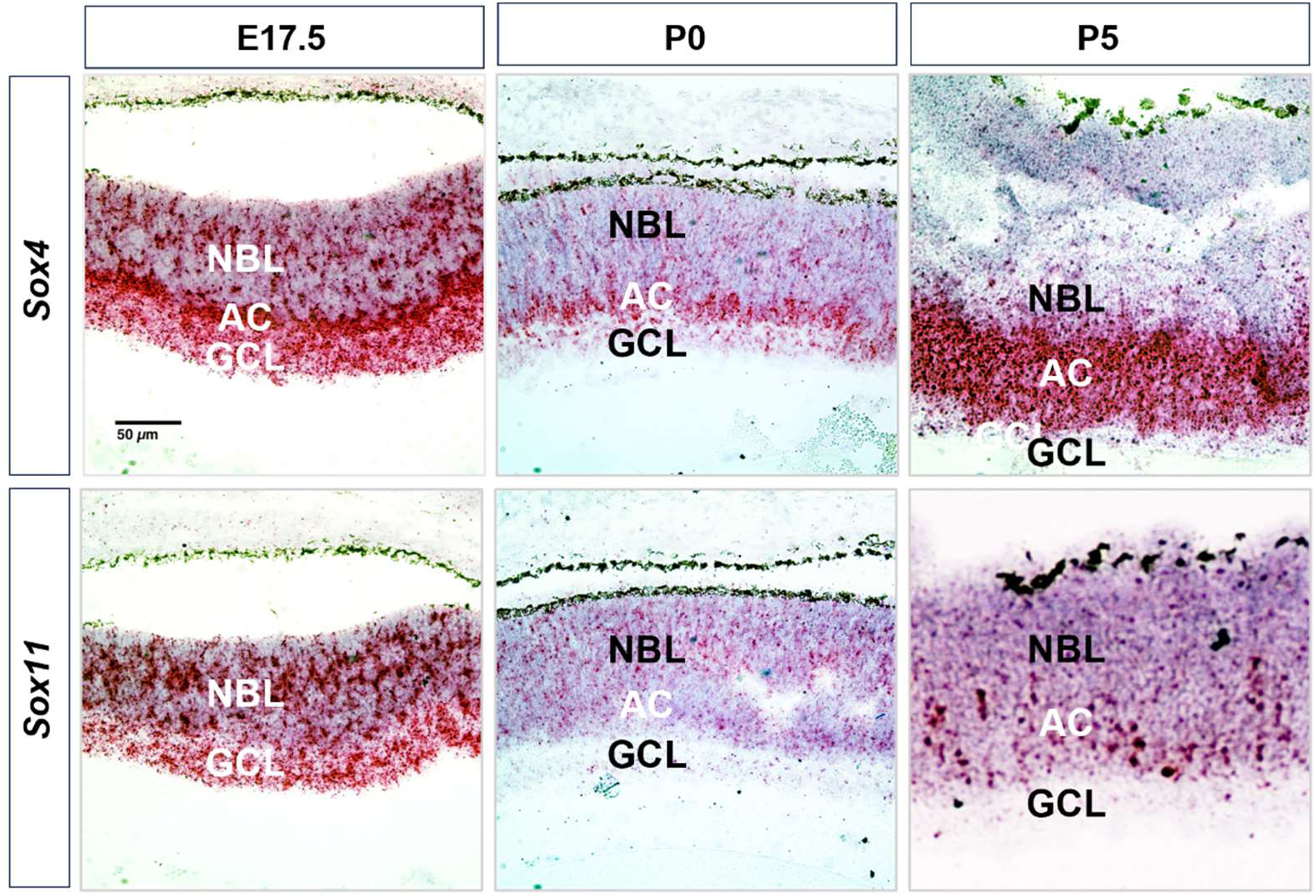
RNAscope in situ hybridization shows that Sox4 and Sox11 continue to be expressed in the retina at later development stages, but with divergent patterns.

**Supplementary Figure 3.**
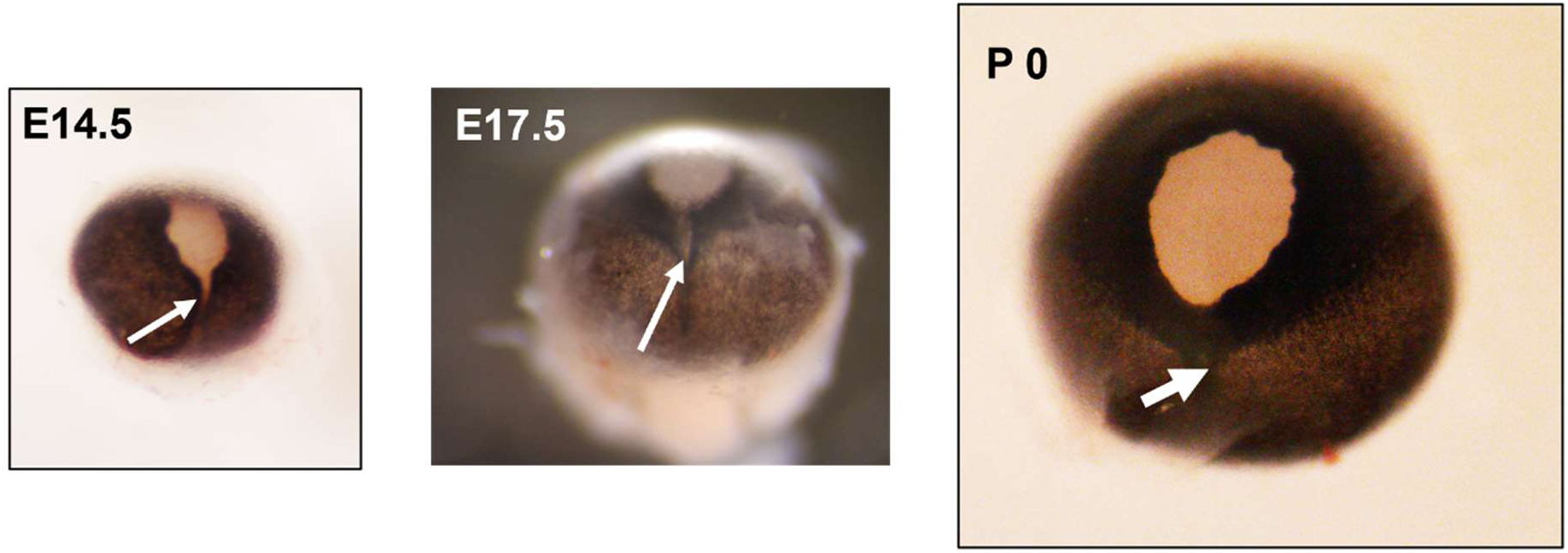
Coloboma manifested in the Sox4/Sox11/Atoh7 tKO retina, which persists in P0.

**Supplementary Figure 4.**
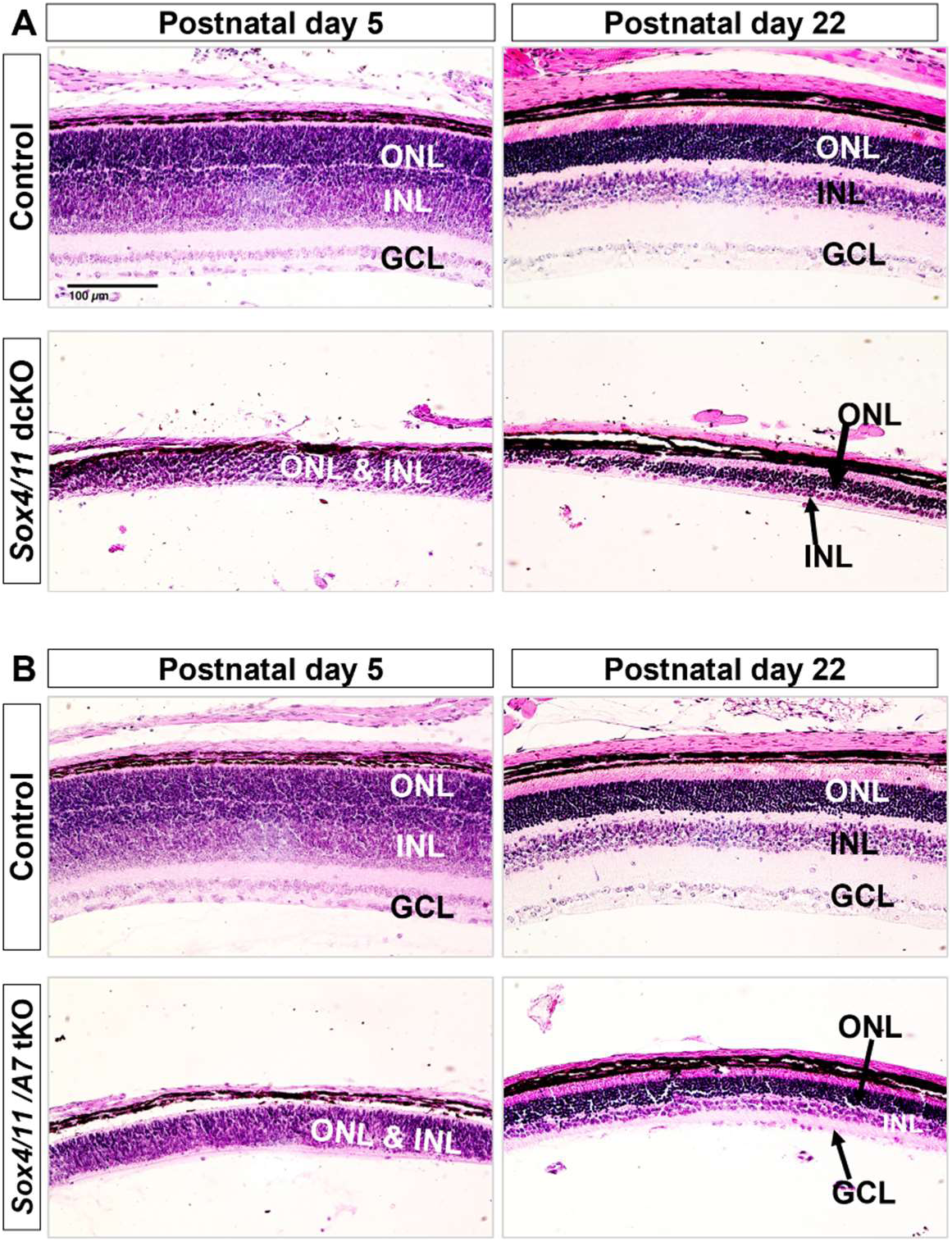
H&E staining of retina sections demonstrating hypoplasia and dysplasia of the dcKO (**A**) and tKO (**B**) retina at P5 and P22.

**Supplementary Figure 5.**
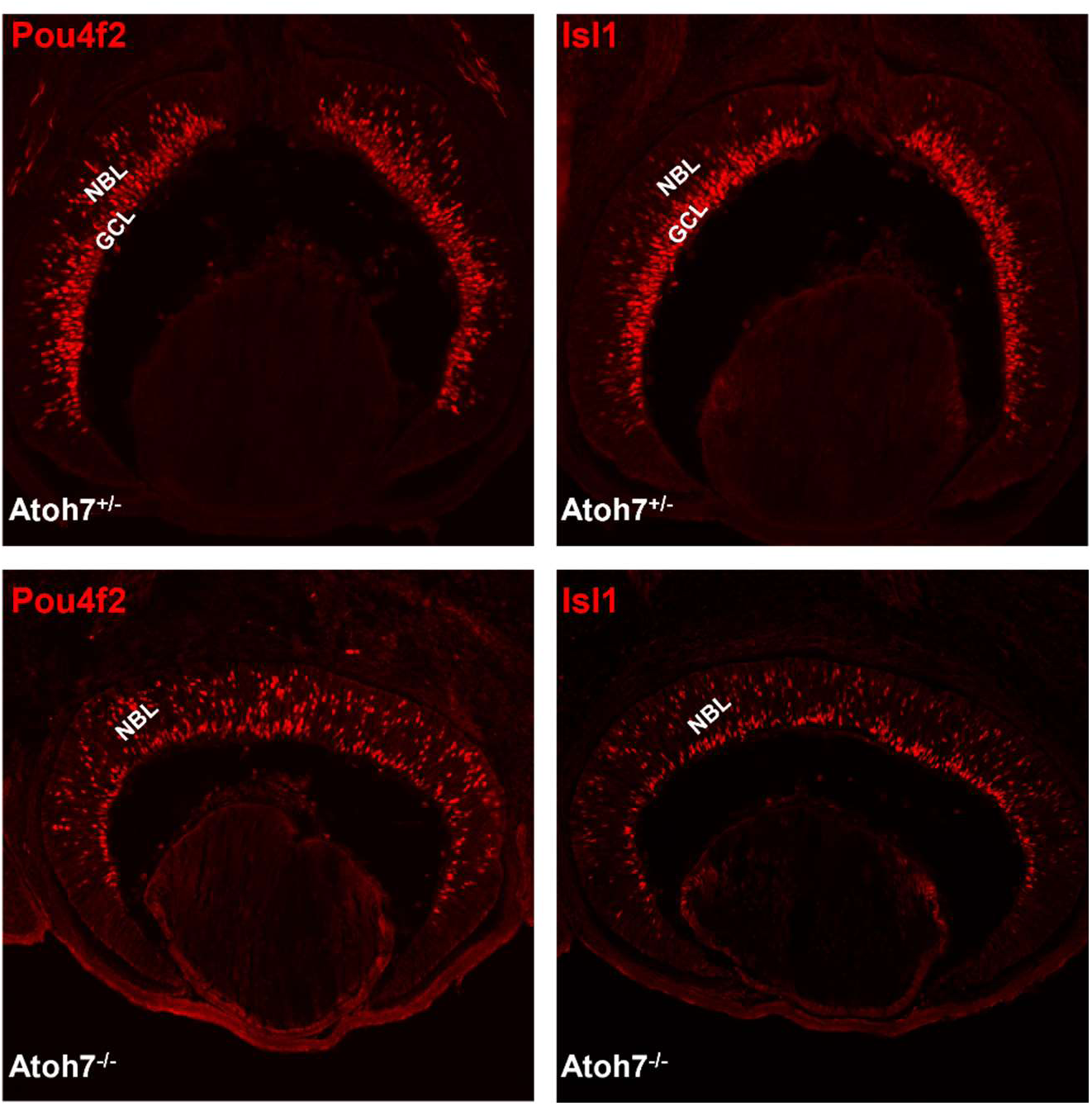
Immunofluorescence staining for Pou4f2 and Isl1 confirms that RGC precursors are still generated in the *Atoh7*-null retina, as indicated by the positive cells in the NBL, but they do not persist, as indicated by the largely diminished GCL.

**Supplementary Figure 6.**
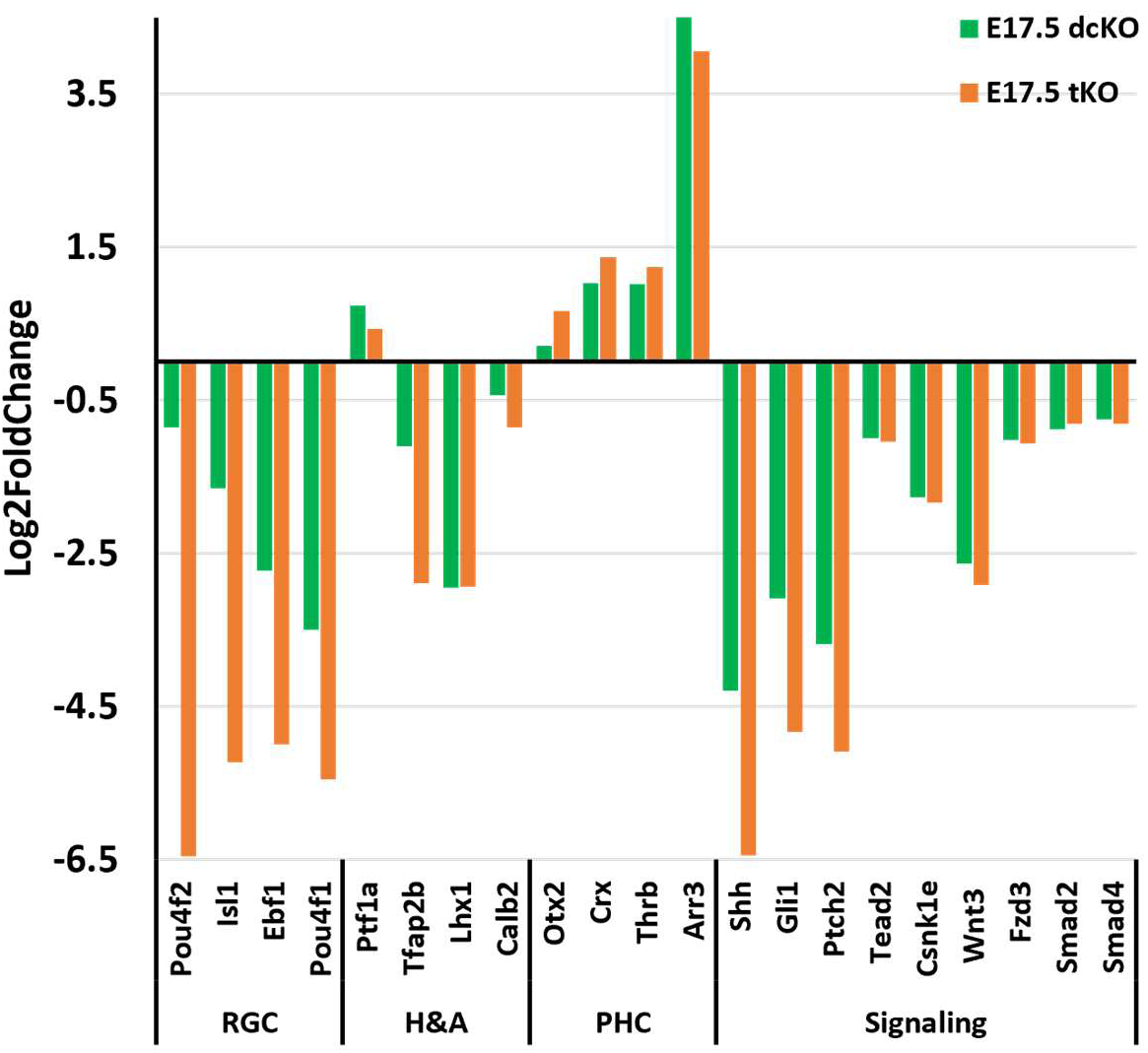
Bar graph displaying different degree of changes of select lineage specific marker genes and genes of signaling pathways in the E17.5 dcKO and tKO retinas.

**Supplementary Figure 7.**
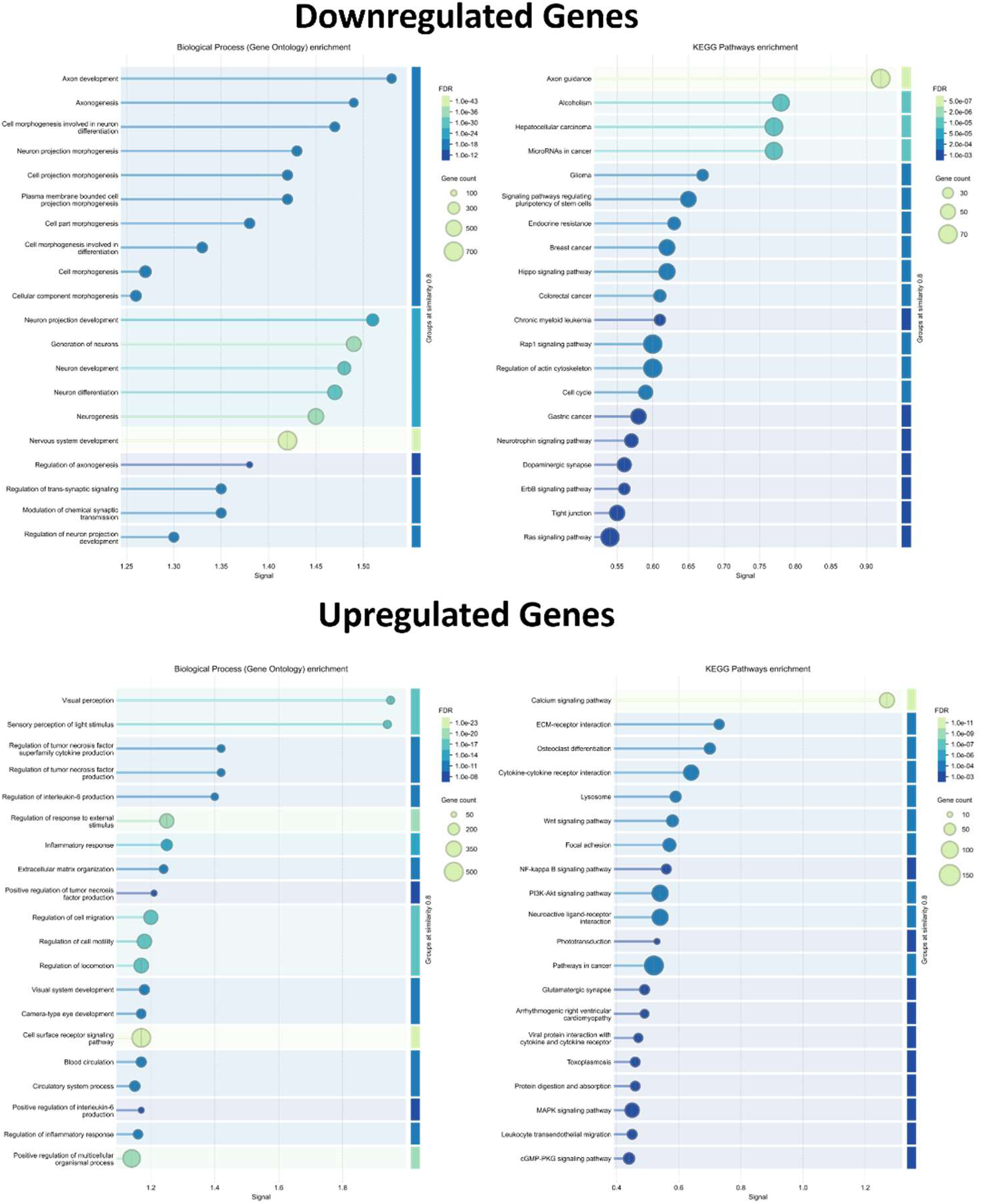
Gene Ontology and KEGG pathway enrichment analysis of down- and up-regulated DEGs in the tKO retina. The dcKO DEGs show similar enrichment but not shown.

**Supplementary Figure 8.**
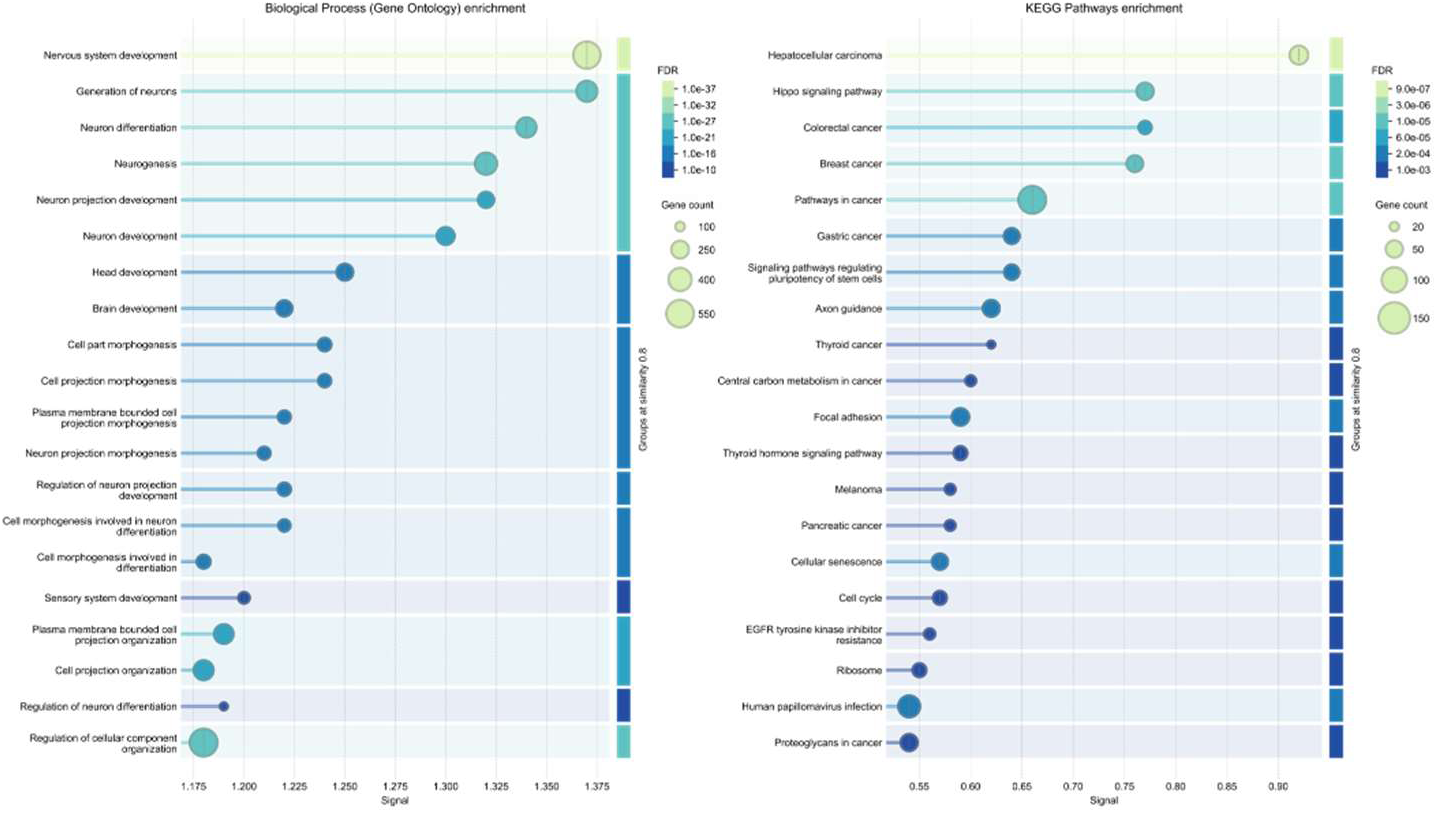
Gene Ontology and KEGG pathway enrichment analysis of Sox11 target genes.

## List of Supplementary tables

**Supplemental Table 1:** DEGs in the E14.5 Sox4/Sox11 dcKO retina.

**Supplemental Table 2:** DEGs in the E17.5 Sox4/Sox11 dcKO retina.

**Supplemental Table 3:** DEGs in the E14.5 Sox4/Sox11/Atoh7 tKO retina.

**Supplemental Table 4:** DEGs in the E17.5 Sox4/Sox11/Atoh7 tKO retina.

**Supplemental Table 5:** Sox11 peaks identified by CUT&Tag and associated genes.

**Supplemental Table 6:** H3K27ac peaks identified by CUT&Tag.

